# MroQ is a Novel Abi-domain Protein That Influences Virulence Gene Expression in *Staphylococcus aureus* via Modulation of Agr Activity

**DOI:** 10.1101/516914

**Authors:** Stephanie Marroquin, Brittney Gimza, Brooke Tomlinson, Michelle Stein, Andrew Frey, Rebecca A. Keogh, Rachel Zapf, Daniel A. Todd, Nadja B. Cech, Ronan K. Carroll, Lindsey N. Shaw

## Abstract

Numerous factors have to date been identified as playing a role in the regulation of Agr activity in *S. aureus*, including transcription factors, antisense RNAs, and host elements. Herein we investigate the product of SAUSA300_1984 (termed MroQ), a transmembrane Abi-domain/M79 protease-family protein, as a novel effector of this system. Using a USA300 *mroQ* mutant we observed a drastic reduction in proteolysis, hemolysis and pigmentation that was fully complementable. This appears to result from diminished *agr* activity, as transcriptional analysis revealed significant decreases in expression of both RNAII and RNAIII in the *mroQ* mutant. Such effects appear to be direct, rather than indirect, as known *agr* effectors demonstrated limited alterations in their activity upon *mroQ* disruption. A comparison of RNA-sequencing datasets for both *mroQ* and *agr* mutants reveal a profound overlap in their regulomes, with the majority of factors affected being known virulence determinants. Importantly, the preponderance of alterations in expression were more striking in the *agr* mutant, indicating that MroQ is necessary, but not sufficient, for Agr function. Mechanism profiling revealed that putative residues for metalloprotease activity within MroQ are required for its Agr controlling effect, however this is not wielded at the level of AgrD processing. Virulence assessment demonstrated that *mroQ* and *agr* mutants both exhibited increased formation of renal abscesses, but decreased skin abscess formation, alongside diminished dermonecrosis. Collectively, we present the characterization of a novel *agr* effector in *S. aureus*, which would appear to be a direct regulator, potentially functioning via interaction with the AgrC histidine kinase.

## Introduction

Pathogenicity in *S. aureus* is both diverse and tightly controlled, being highly adaptive to the surrounding environment and available resources (1). In order to survive and persist within the host, this pathogen produces a wide array of virulence factors throughout the different stages of infection (2). For example, during earlier phases of growth, adhesins and other binding proteins are actively synthesized (e.g. Spa, FnbAB), which play a role in colonization (2, 3). In contrast, in later phases, these surface proteins are repressed, whilst secreted toxins, proteases, and exoenzymes are produced to facilitate invasion and dissemination (2, 4). This coordination is facilitated by an array of global regulators of gene expression (2, 5–10).

The *agr* quorum sensing system is one of the most important regulators of virulence in *S. aureus* (11). It is a quorum sensing system comprised of two transcriptional units, RNAII and RNAIII, each controlled by its own promoter, P2 and P3, respectively. RNAII encodes the *agr* operon, which is comprised of four genes, *agrBDCA* (12). The quorum sensing system functions in three parts: i) a quorum sensing module comprised of AgrB and AgrD; ii) a two-component system (TCS) AgrC and AgrA; iii) an effector molecule, termed RNAIII, which serves as a regulatory RNA (2, 12, 13). The pheromone for the *agr* system is known as autoinducing peptide (AIP), and is a derivative of the AgrD polypeptide. In order to reach its active form (AIP), AgrD is initially processed by AgrB, a multifunctional endopeptidase that also acts as a chaperone, followed by processing by the SpsB signal peptidase (14, 15). The AIP peptide is exported by AgrB, and accumulates extracellularly until a threshold is reached, when it binds AgrC, the histidine kinase (HK) of the TCS.

This leads to autophosphorylation of the protein, (12, 16) followed by transfer of that phosphate to AgrA, the cytoplasmic response regulator (RR) (15). Upon activation, AgrA directly binds to the P2 and P3 promoters, upregulating expression of RNAII, as a positive feedback loop, as well as RNAIII (12). RNAIII subsequently modulates the expression of an assortment of genes involved in virulence, thus exerting the crucial role of *agr* in *S. aureus* pathogenesis. To date, numerous regulators of *agr* have been identified (5, 7, 8, 17–21), including transcription factors, such as SarR and CodY (5, 20), antisense RNAs such as the psm-mec RNA transcript (22), and host factors such as the serum protein apolipoprotein B (23).

Abi proteins have previously been noted to play a protective role against viral infection by halting cellular activity in infected bacterial cells, so as to prevent further replication of the invading bacteriophage (24). These proteins are also members of the M79 metalloprotease family, which in eukaryotes are also noted as CaaX prenyl proteases (25, 26). These eukaryotic proteases are involved in post-translational modification through a mechanism known as prenylation, where the ‘aaX’ motif of target proteins are cleaved from the C-terminus by M79 enzymes, followed by the addition of an isoprenoid on the remaining cysteine residue (27). Work by our group exploring the prenylation process in bacteria has demonstrated that the C-terminal CaaX motif is present in only a handful of proteins, suggesting that this mechanism is not conserved in prokaryotes (28). Indeed, only a single protein has been shown to be prenylated in bacteria: ComX from Bacillal species is known to be prenylated at a conserved C-terminal tryptophan residue (29), although it is not known if proteolysis is a requirement for this event.

Beyond a role in protecting against phage infection, Abi proteins have been characterized in only two other studies. The first, Abx1, has been the subject of study in Group B Streptococci (27). Notably, an *abx1* mutant demonstrated a significant decrease in hemolytic activity and pigment production, which was mediated via increased CovSR activity (27). In this work, Abx1 was shown to positively regulate CovS, a histidine kinase involved in virulence gene regulation, by forming a signaling complex. Moreover, the interaction was established to be direct, via the two CovS transmembrane domains, but not dependent on Abx1 proteolytic activity (27). More recently, an Abi-domain protein in *S. aureus*, SpdC, was characterized and shown to interact with the histidine kinase of the WalKR two-component system, which is involved in controlling cell wall metabolism (30). SpdC was also shown to interact with other histidine kinases, including SaeS, ArlS, and VraS, through interaction between transmembrane domains (30). Of note, although an Abi protein, SpdC lacks the conserved catalytic residues of M79 enzymes, and is thus a non-peptidase homolog. As such, although still relatively underexplored, it is becoming increasingly apparent that Abi proteins in prokaryotes possess important regulatory functions that are seemingly mediated through interaction with the histidine kinase modules of two-component systems (27, 30).

In *S. aureus*, beyond SpdC, there are 5 other Abi-domain proteins encoded within the genome (**Supplemental Figure S1**). Herein, we investigate the function of one of these, SAUSA300_1984, located upstream of the *agr* operon, which we name MroQ. We show that mutation of *mroQ* results in decreased expression and abundance of secreted proteases, hemolysins, and toxins, as well increased production of surface associated proteins. These effects were shown to be *agr* dependent, with the *mroQ* mutant demonstrating decreased expression of both RNAII and RNAIII. Importantly, known regulators of *agr* demonstrated limited alterations in transcription, potentially indicating a direct role for MroQ in controlling *agr* activity. We also noted decreased pigment production and enhanced biofilm formation in the *mroQ* mutant, which mirrored that of an *agr* mutant. Finally, we observed decreased bacterial burden and dermonecrosis for *mroQ* mutant infected mice using a model of skin abscess formation. Collectively, we present the characterization of a novel *agr* effector in *S. aureus*, which would appear to be a direct regulator, potentially functioning via interaction with the AgrC HK component.

## Results

### MroQ serves to control protease production in *S. aureus*

Given the longstanding interest of our laboratory in *S. aureus* secreted proteases, we screened the Nebraska Transposon Mutant Library (31) for novel factors that could regulate their production. Using this mutant collection and casein agar plates, we identified an insertion in gene SAUSA300_1984 (hereafter named *mroQ*, <u>M</u>embrane Protease <u>R</u>egulator <u>o</u>f Agr <u>Q</u>uorum Sensing) that elicited a profound reduction in secreted protease activity (**Fig 1A**). To explore this more broadly, we next analyzed proteolytic activity through gelatin zymography (**Fig 1B**). Here, the *mroQ* mutant demonstrated a substantial decrease in levels of multiple secreted proteases, which was rescued upon complementation *in trans;* indeed, complementation with a multi-copy plasmid actually led to an increase in the production of secreted proteases in comparison to our wild type strain. Next, to determine if the influence of MroQ on protease production was a consequence of modulated activity or altered transcription, we used a reporter gene fusion for the protease aureolysin. When this fusion was transduced into the USA300 wild type and *mroQ* mutant we observed a significant impairment in aureolysin transcription in the mutant strain (**Fig 1C**), thus indicating the effects are wielded at the level of gene expression.

**Figure 1.**
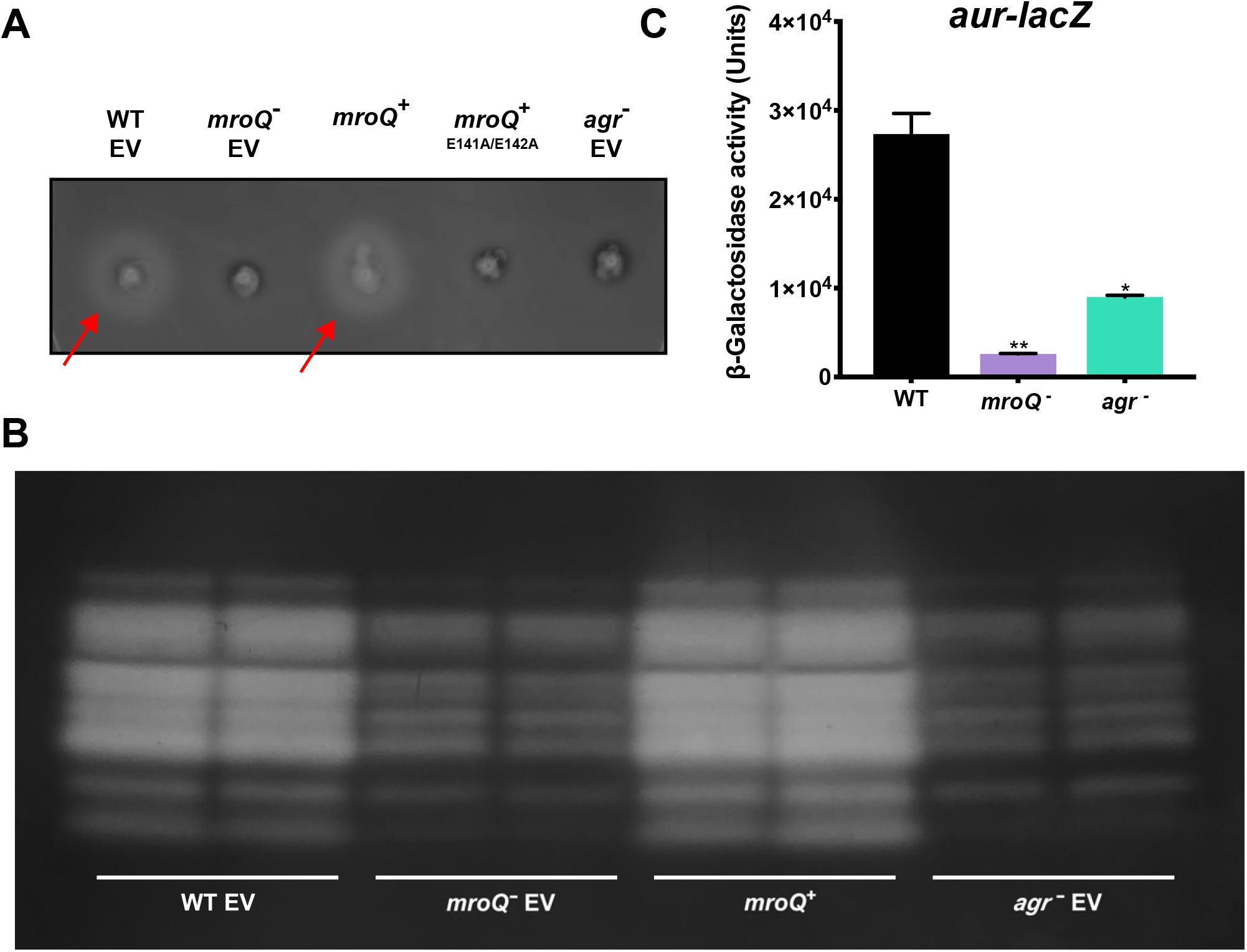
Disruption of *mroQ* Results in Diminished Proteolytic Activity in *S. aureus*. The wild type, *mroQ* mutant, *mroQ* complemented strain, *mroQ* site-directed mutant (E141A/E142A) complemented strain and *agr* mutant strain were plated on casein nutrient agar to observe changes in proteolysis. Red arrows highlight zones of proteolysis. (**A**). Gelatin zymography of indicated strains. Note: White zones of clearing indicated proteolysis of gelatin substrate. (**B**). Activity of an *aur-lacZ* fusion in the wild type, *mroQ* mutant, and *agr* mutant strains. Data is from 4h of growth, which is a peak window of aureolysin expression (**C**). Student’s t-test was used to determine statistical significance, *= p<0.05, **=p< 0.01. Error bars are shown with SEM. EV = Empty Vector.

### MroQ regulates the production of a wealth of *S. aureus* virulence determinants

To determine if the observed effects were specific to proteases, we next explored whether the production of other virulence factors was influenced by MroQ. Using a blood agar plate, we observed a significant decrease in hemolytic activity for the mutant strain when compared to the wild type (**Fig 2A**) that was restored upon complementation *in trans*. This activity was explored quantitatively using a hemolysin assay with whole human blood, revealing a substantial decrease in the ability of the mutant to lyse human erythrocytes; which was again complementable (**Fig 2B**). To assess whether this effect was mediated at the transcriptional level, we used a reporter gene fusion for the primary hemolysin in *S. aureus, hla*. When the activity of this construct was assessed between the wild type and *mroQ* null strain we observed a significant decrease in the mutant, indicating these effects are also mediated at the level of gene expression (**Fig 2C**). To further investigate the role of MroQ, and its influence on other virulence determinants, we performed western blots for the Panton Valentine Leukocidin (PVL) toxin and the surface exposed virulence factor, Surface Protein A (Spa). Upon analysis we determined that PVL was highly abundant in the wild type and *mroQ* complement strain, however was entirely absent in the *mroQ* mutant (**Fig 2D**). Conversely, for blots with Spa we found increased abundance in the *mroQ* mutant, whereas we noted an absence in the wild type and *mroQ* complement strain (**Fig 2E**). To ensure that these substantial changes in virulence factor production were not the result of a simple growth defect in the mutant, we measured cell density over time. We found no significant difference when performing these studies between the wild type, mutant and complemented strains (data not shown).

**Figure 2.**
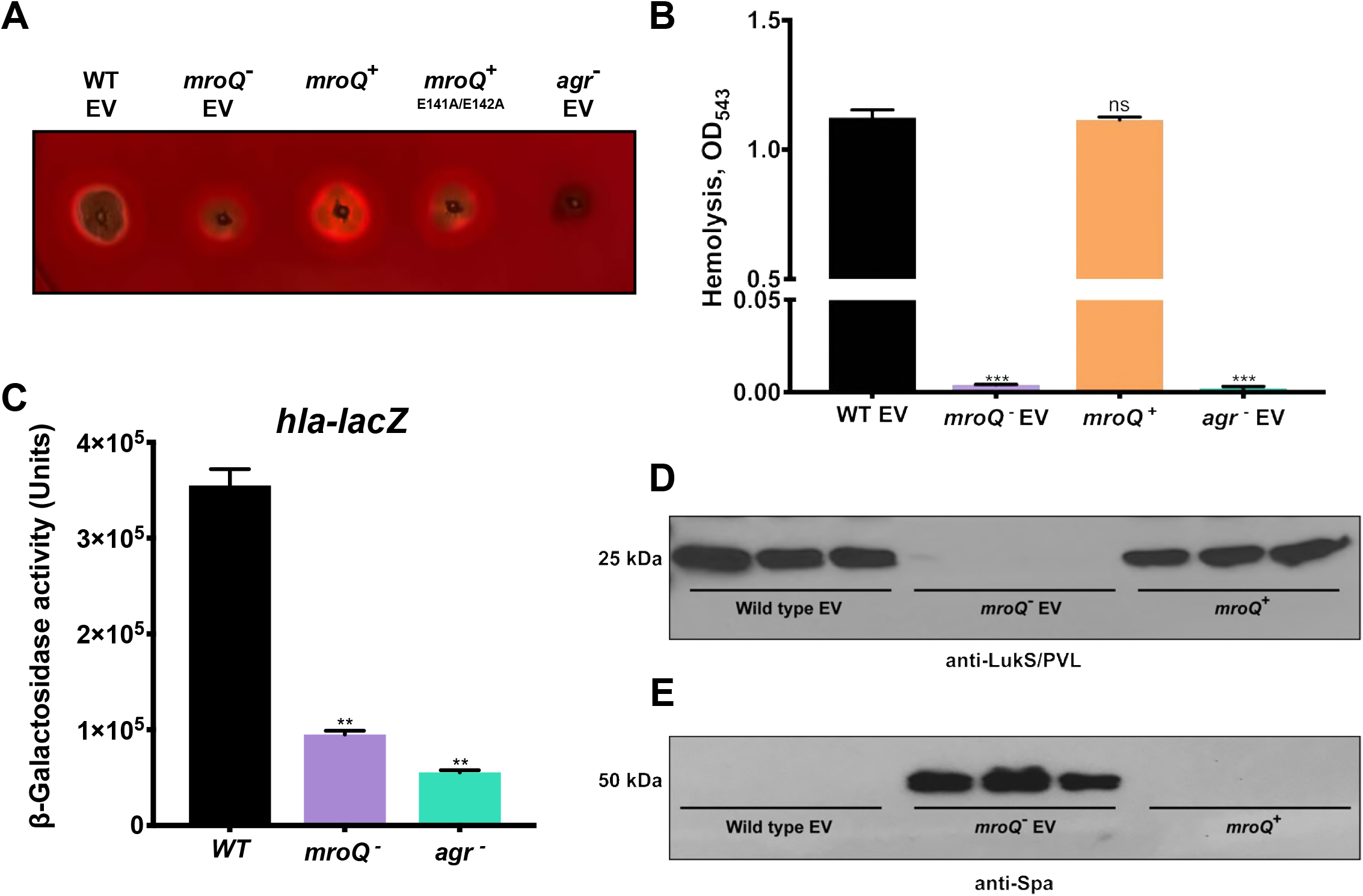
MroQ Globally Regulates Virulence Factor Production in *S. aureus*. The wild type, *mroQ* mutant, *mroQ* complemented strain, *mroQ* site-directed mutant (E141A/E142A) complemented strain and *agr* mutant strain were plated on sheep blood agar to observe changes in hemolysis (**A**). Hemolytic activity was measured via lysis of erythrocytes in whole human blood (**B**). Activity of a *hla-lacZ* fusion in the wild type, *mroQ* mutant, and *agr* mutant strains. Data is from 4h of growth, which is a peak window of α-hemolysin expression (**C**). Immunoblots for abundance of the LukS component of the Panton Valentine Leukocidin (PVL, **D**), and Surface Protein A (SpA, **E**) in the wild type, *mroQ* mutant and complemented strains. Student’s t-test was used to determine statistical significance, **= p<0.01; ***= p<0.001. Error bars are shown with SEM. EV = Empty Vector.

### MroQ mediates its effects via modulation of *agr* activity in *S. aureus*

Our findings thus far for the *mroQ* mutant are in line with that which one might expect for a strain exhibiting altered activity of the central virulence factor regulator, *agr*. Indeed, when we repeated our proteolysis and hemolysis activity and transcription studies in Figures 1 and 2 with an *agr* mutant strain, we saw very similar results to the *mroQ* mutant. As such, we next sought to assess changes in promoter activity for the *agr* P2 and P3 promoters using qPCR analysis (**Fig 3**). When abundance of the RNAII (**A**) and RNAIII (**B**) transcripts was measured we observed a marked decrease for both elements in the mutant as compared to the parent. Specifically, activity from the P2 promoter was decreased by 15.6-fold, whilst activity from P3 was decreased by a striking 292-fold.

**Figure 3.**
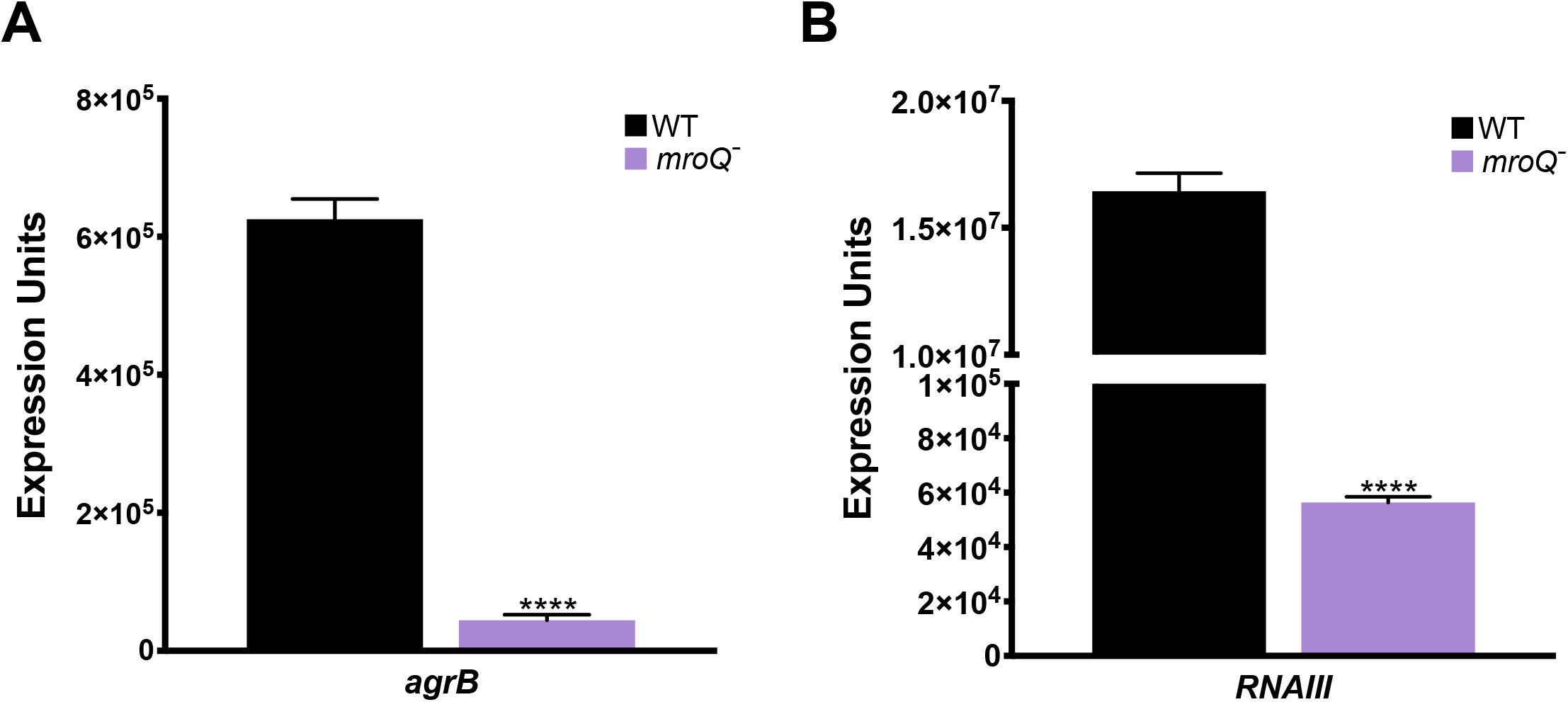
*RNAII* and *RNAIII* are Significantly Downregulated Upon *mroQ* Disruption. qPCR was performed on the *mroQ* mutant and USA300 wild type strains grown in TSB to late-exponential phase. Target expression levels were normalized to 16s rRNA. *agrB* was used as a measure of *RNAII* transcript abundance. Student’s t-test was used to determine statistical significance, **** = p ≤ 0.0001. Error bars are shown with SEM.

To further clarify these effects, we next assessed transcription of a range of elements downstream of Agr, whose transcription is dependent on its activity. Here, some very notable changes in expression between the wild type and mutant were observed, including numerous genes involved in virulence that are positively regulated by *agr*. Specifically, decreases in the mutant strain were observed for: *psmα1* (2666-fold), *sspA* (57.3-fold), *lukS-PV* (35.4-fold), and *lip* (132.2-fold). Further, the notable regulator *saeR*, which is positively influenced by the *agr* system, demonstrated a 3.8-fold decrease in transcription in the mutant. Additionally, *spa*, which is repressed by the *agr* effector RNAIII, demonstrated a 792-fold increase in transcription in the mutant, as did the notable SarA homolog, *sarS* (which is also negatively regulated by *agr)*, which was found to have a 2.1-fold increase in transcription. Another surface protein that is repressed by RNAIII, *sdrD*, also demonstrated a 2.7-fold increase in transcription in the mutant strain (**Fig 4**).

**Figure 4.**
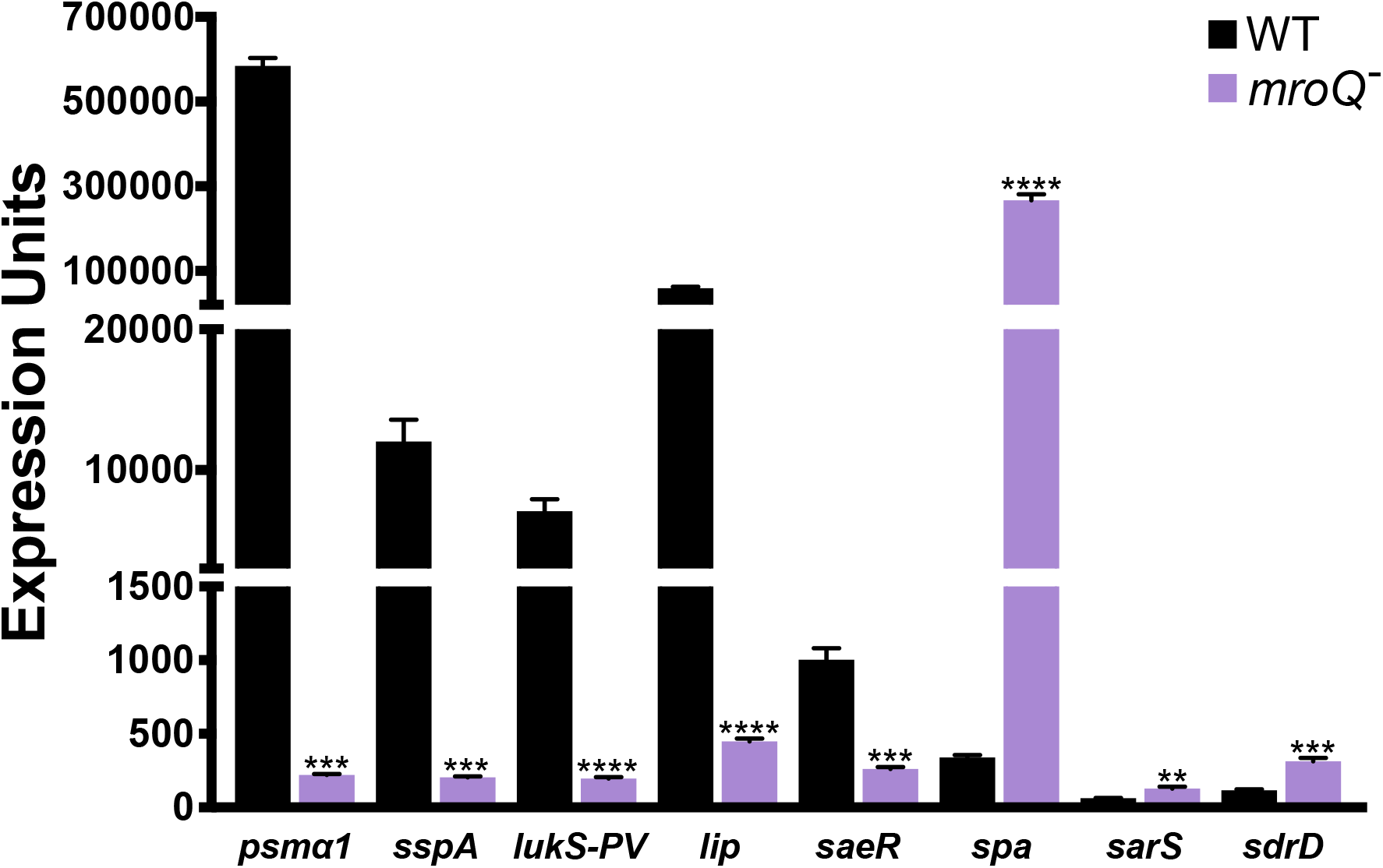
The Expression of Genes Regulated by Agr are Altered in the *mroQ* Mutant. qPCR was performed on the *mroQ* mutant and USA300 wild type strains grown in TSB to late-exponential phase. Expression levels were normalized to 16s rRNA. Student’s t-test was used to determine statistical significance, **=p< 0.01, *** = p ≤ 0.001, **** = p ≤ 0.0001. Error bars are shown with SEM.

To determine if these were likely direct or indirect effects, we next sought to measure the expression of a cadre of known upstream effectors of the *agr* system. Accordingly, expression of positive (*sarA, sarZ, sarU*, and *mgrA*) and negative (*arlR, sarR, sarX, rsr, codY*, and *sigB*) regulators of *agr* activity were studied by qPCR in the *mroQ* mutant and parental strain (**Fig 5, Supplemental Table S1**). Importantly, when these experiments were performed, we observed no significant alteration in the transcription of key *agr* repressors, such as *sigB* (1.1-fold decrease), *sarX* (1.1-fold decrease), *rsr* (1.2-fold decrease), and *codY* (1.2-fold increase). Moreover, although changes of merit were observed for two other *agr* repressors, *arlR* (1.8-fold decrease) and *sarR* (8.2-fold decrease), these were both negative effects, and thus do not explain the phenotypes observed. When looking at positive regulators of *agr*, we did observe decreased expression for a number of factors, but none at levels that would prove causative to the large decreases in *agr* activity (292-fold decrease in P3 activity in the *mroQ* mutant) thus far observed: *sarA* (3.1-fold decrease), *sarZ* (2-fold decrease), *sarU* (2.1-fold decrease), and *mgrA* (1.9-fold decrease). As such, although not definitive, it would appear that the positive impact of MroQ on *agr* activity is either mediated by direct interaction, or via an as yet unknown regulator of this system.

**Figure 5.**
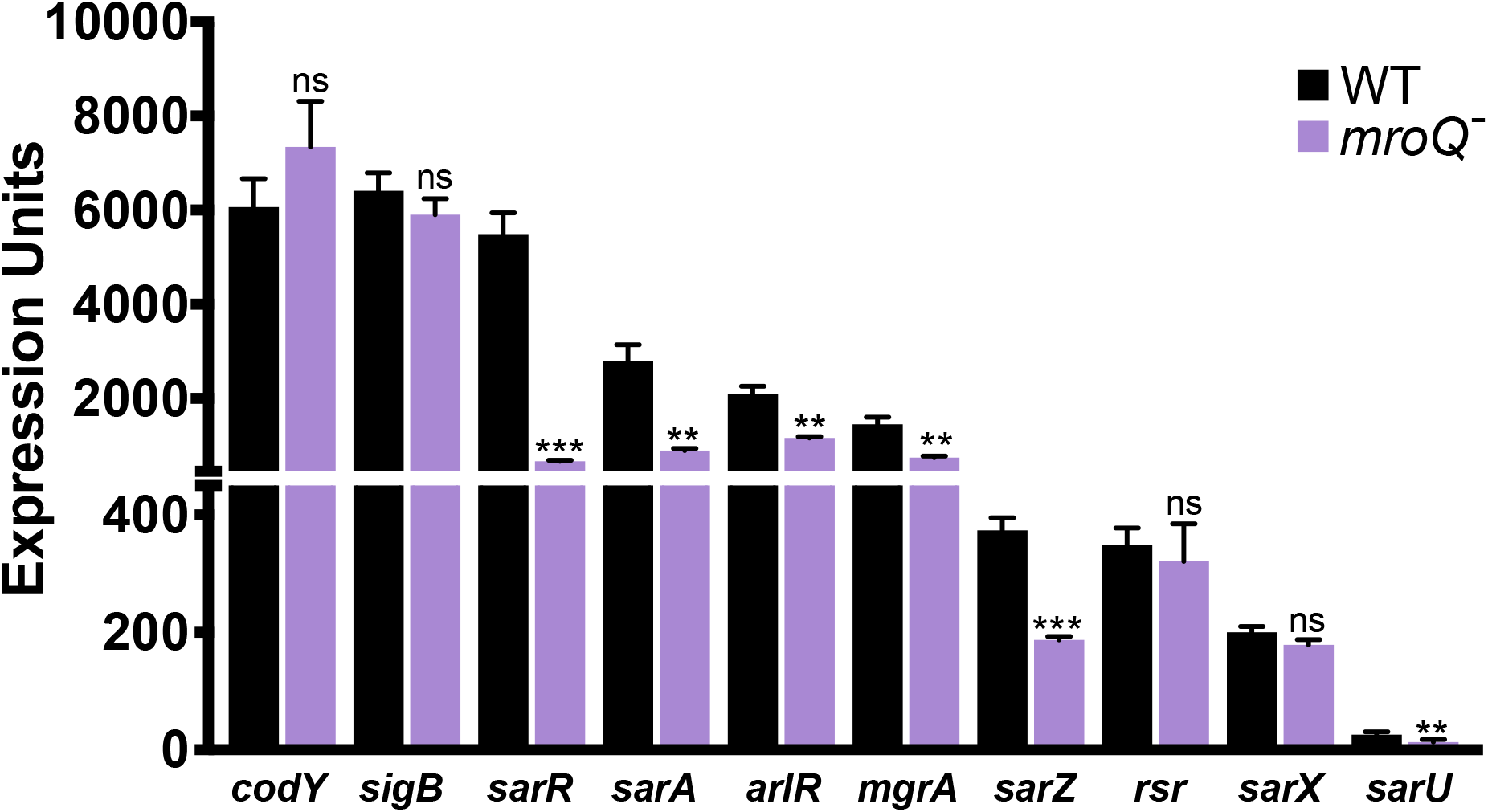
Alterations in Expression of Known Regulators of *agr* in the *mroQ* Mutant. qPCR was performed on the *mroQ* mutant and USA300 wild type strains grown in TSB to late-exponential phase. Target expression levels were normalized to 16s rRNA. Student’s t-test was used to determine statistical significance, ** = p ≤ 0.01, *** = p ≤ 0.001, n.s. = not significant. Error bars are shown with SEM.

### Probing the role of MroQ using a global transcriptomic approach

To gain a more global insight into the regulatory effects of MroQ in *S. aureus*, we performed RNA-seq analysis on our USA300 *mroQ* mutant and its parental strain. Strains were grown in TSB until late-exponential phase, and RNA was isolated for sequencing. A genomic map was created to demonstrate the changes in transcriptome between the *mroQ* mutant and the parent, as well as fold changes in gene expression denoted by a heat map (**Fig 6A**). Upon analysis, we found that expression of 123 genes/sRNAs was decreased in the *mroQ* mutant, and 109 were increased at a level of ≥3-fold (**Fig 6B, Supplemental Table S2**). Importantly, these studies confirmed our previous data, demonstrating that all components of the Agr system were indeed downregulated in the *mroQ* mutant, with the largest decrease observed in its two-component system, AgrC (21.6-fold) and AgrA (22.2-fold). When changes were clustered ontologically based on known or predicted function, the most downregulated group was virulence factors, with 28 genes decreased in expression in the mutant strain (**Fig 6C**). Furthermore, our previous data regarding virulence factors was supported with very significant changes in expression for genes encoding proteases, toxins, and surface-associated proteins. For example, PSM expression showed a 695-fold decrease, and the *lukS-PV* toxin demonstrated a 63.5-fold decrease in the mutant. Notable proteases such as *splA*, the first gene in the *spl* operon, demonstrated a 903-fold downregulation, *sspA* demonstrated a 47-fold decrease, and *scpA* a 10-fold decrease in the mutant strain. When looking at the MSCRAMMs, *sdrD* and *spa* demonstrated a 5.2-fold and 58.1-fold increase in transcription, respectively, in our *mroQ* mutant.

**Figure 6.**
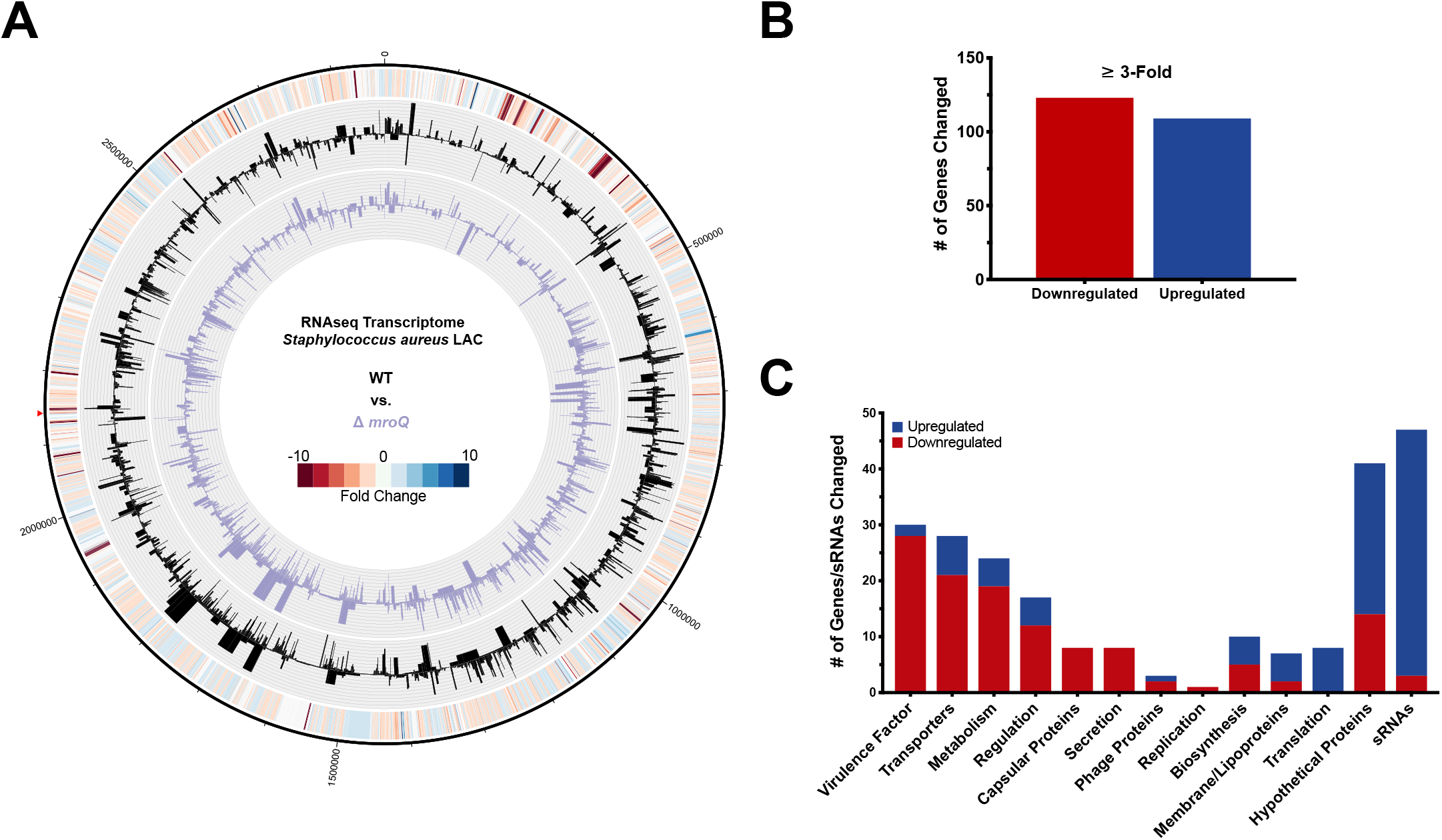
*mroQ* Disruption Produces a Global Alteration in *S. aureus* Gene Expression. RNA-seq was performed on the USA300 wild type and its *mroQ* mutant strain. Cultures were grown in TSB to late-exponential phase in biological triplicate. RNA was isolated, and the total RNA of replicates was processed for sequencing on an IonTorrent PGM. A genomic map was created to denote changes in the *mroQ* mutant transcriptome (inner circle, purple), compared to the wild type (middle circle, black). The outermost circle is a heat map demonstrating fold changes in expression (**A**). Depiction of the number of genes upregulated and downregulated (**B**), which were sorted into ontological groups based on known or predicted function (**C**).

Considering the clear evidence indicating decreased Agr activity in the *mroQ* mutant, and that the most profound changes observed in transcription are for virulence determinants, we next decided to examine a full *agr* operon knockout using RNA-seq analysis. This strain, alongside its parent, was grown in TSB until late-exponential phase, and RNA was isolated for sequencing. A genomic map was created to demonstrate the changes in transcriptome between the *agr* mutant and wild-type, as well as fold changes in gene expression via a heat map (**Fig 7A**). Implementation of a 3-fold cut off demonstrated that the expression of 222 genes/sRNAs was decreased, and 262 were increased in the *agr* null strain (**Fig 7B, Supplemental Table S3**). When reviewed ontologically, the cluster of genes that appeared to be most downregulated was again virulence factors (**Fig 7C**). Upon comparing the *mroQ* mutant transcriptome to that of the *agr* mutant, we found 55% similarity in altered genes and sRNAs (**Supplemental Figure S2**). Of these 129 shared genes, a quarter of them encode for virulence factors. It is important to note that the percentage of shared genes/sRNAs is likely higher between the two mutants, as numerous genes identified in the *mroQ* mutant dataset are known to be regulated by Agr, however due to parameters implemented during analysis, were not present in the *agr* mutant dataset due to being too lowly expressed upon deletion of *agr*. To validate findings for both transcriptome studies, we performed qPCR analysis on a random selection of genes (**Supplemental Table S4**), with each showing changes that reproduce that of our RNA-seq datasets.

**Figure 7.**
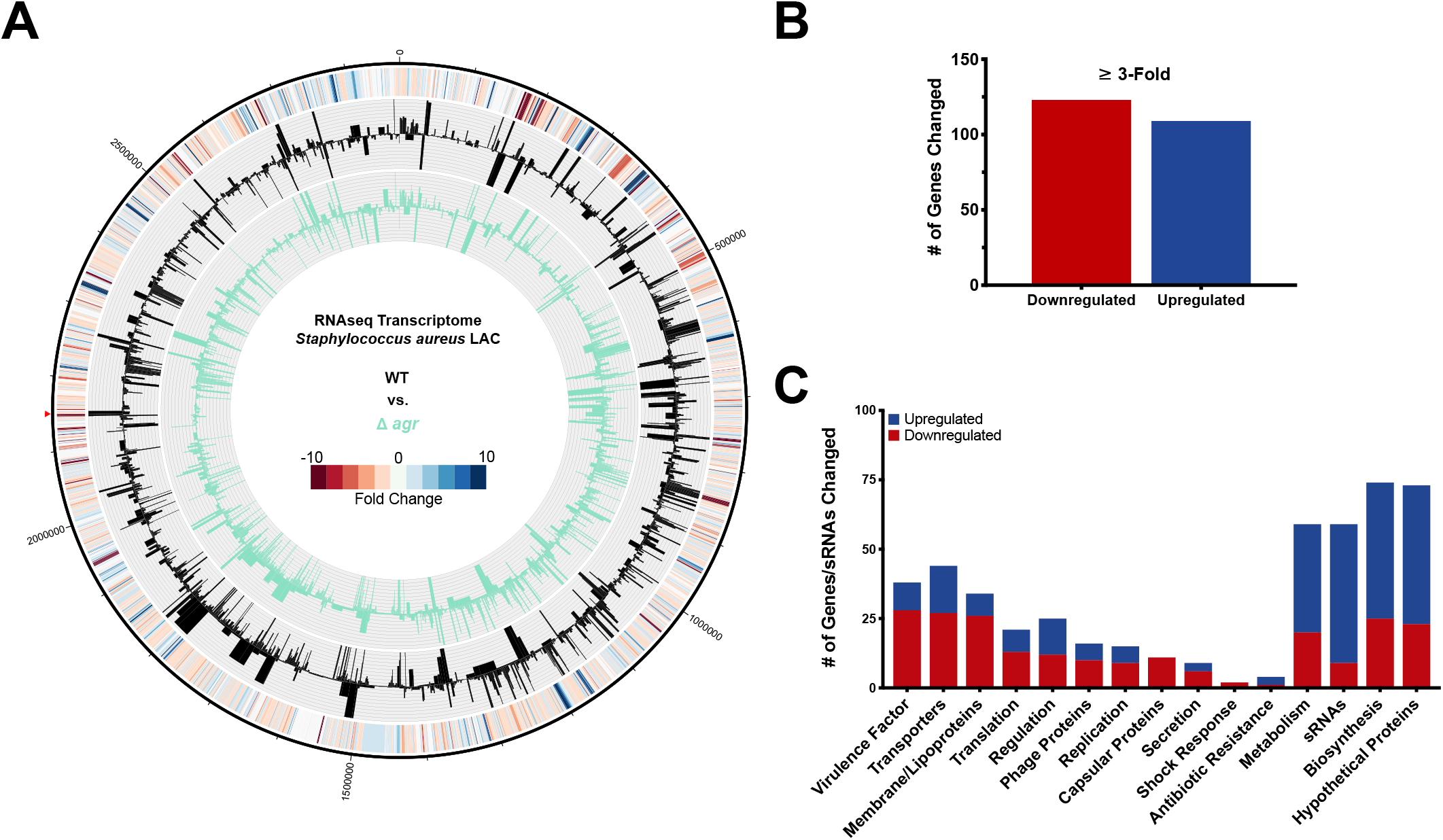
Transcriptomic Profiling of an *agr* Operon Deletion Mutant. RNA-seq was performed on the USA300 wild type and its *agr* mutant strain. Cultures were grown in TSB to late-exponential phase in biological triplicate. RNA was isolated, and the total RNA of replicates was processed for sequencing on an IonTorrent PGM. A genomic map was created to denote changes in the *agr* transcriptome (inner circle, green), compared to the wild type (middle circle, black). The outermost circle is a heat map demonstrating fold changes in expression (**A**). Depiction of the number of genes upregulated and downregulated (**B**), which were sorted into ontological groups based on known or predicted function (**C**).

Collectively, this profound similarity in transcriptomes again suggests that mutation of *mroQ* elicits an *agr* mutant-like phenotype. It is important to note, however, that ablated MroQ activity elicits an intermediate *agr* deficient phenotype rather than a complete null effect. This is evidenced through a comparison of fold changes for primary virulence factors between the *mroQ* and *agr* mutants (**Table 3**), where *mroQ* disruption results in less profound changes in expression than *agr* deletion.

### Mutation of *mroQ* enhances biofilm formation

Given the noted increased in abundance of surface proteins and the marked decrease in virulence factor activity in our *mroQ* mutant strain, we hypothesized that *mroQ* disruption would lead to enhanced biofilm formation, as seen with *agr* mutant strains (32). Using the classic crystal violet assay for biofilm formation, described by ourselves and others previously (33), we determined that the *mroQ* mutant strain did indeed produce a more robust biofilm than our USA300 wild type strain, with levels similar to that of an *agr* mutant (**Fig 8**). This effect was restored to wild type biofilm formation levels following complementation with *mroQ in trans*. These results are consistent with those previously seen for *agr* mutants in multiple different *S. aureus* backgrounds (32, 34).

**Figure 8.**
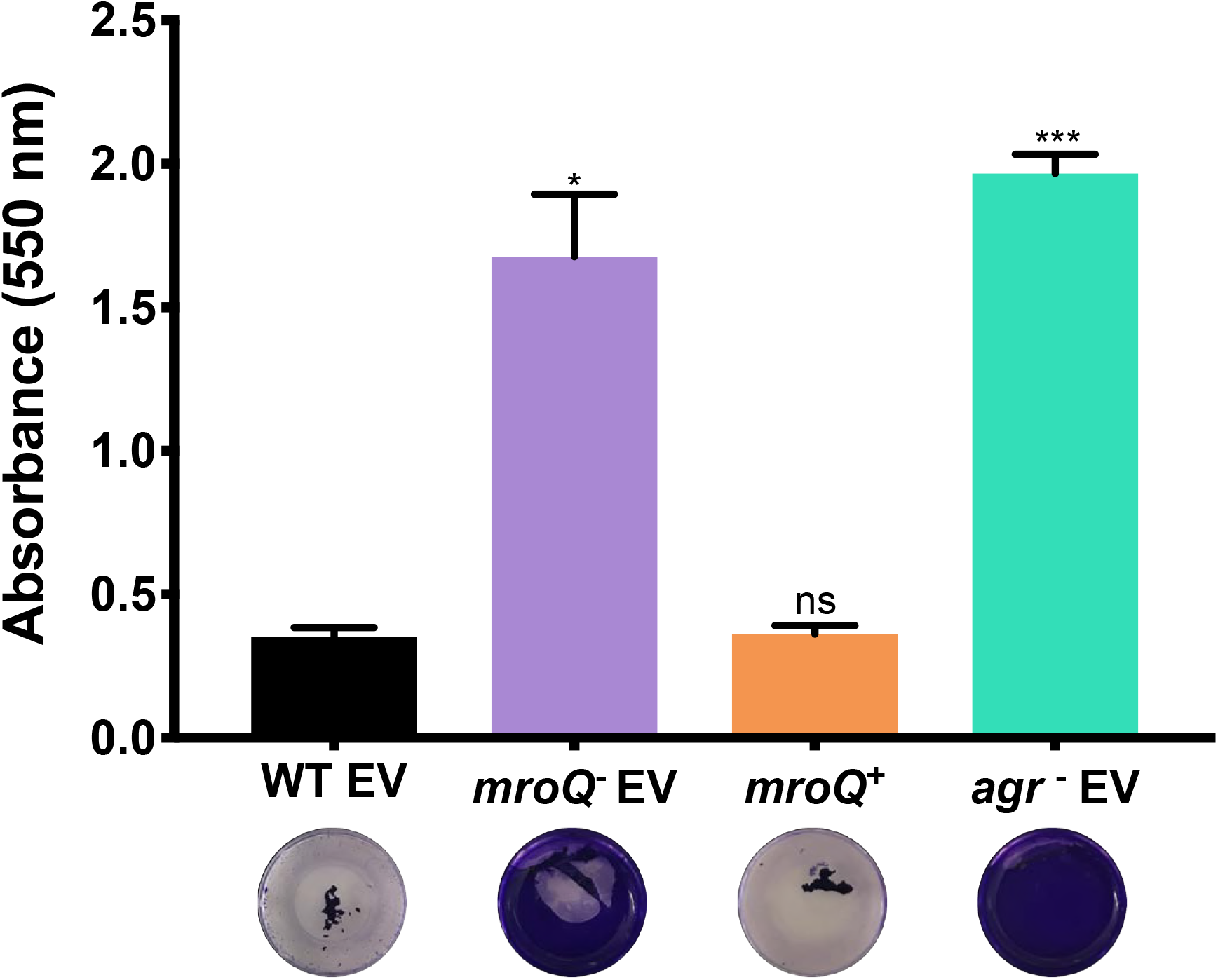
The *mroQ* Mutant Demonstrates an Increased Capacity for Biofilm Formation. Strains were grown in 24-well plates in TSB media containing 5% human plasma. Biofilm formation was quantified with crystal violet staining, followed by extraction with 100% ethanol and OD_550_ measurement. Assays were performed in biological triplicate and technical duplicate. Student’s t-test was used to determine statistical significance, *= p<0.05, *** = p ≤ 0.001. Error bars shown with SEM. EV = Empty Vector, ns = not significant.

### Disruption of *mroQ* leads to an *agr*-mediated decrease in staphyloxanthin production and impaired tolerance to oxidative stress

During our work with the *mroQ* and *agr* mutant strains we observed that both appear to have a decrease in the characteristic golden pigment produced by *S. aureus* (**Fig 9A**). To determine if this was mediated at the level of gene expression we performed qPCR on one of the carotenoid biosynthesis operon genes, *crtM*, for both the *mroQ* and *agr* mutants, revealing no significant difference in gene expression (data not shown). Next, we examined *ispA* transcription, which has been previously shown by our lab to have a significant role in production of farnesyl pyrophosphates (FPPs), the starting product in the staphyloxanthin biosynthesis pathway (28, 35). Here, we once more found no significant difference in gene expression (data not shown). Of note, our qPCR analysis previously demonstrated no significant changes in *sigB* expression, which is absolutely required for expression of the *crt* operon, in the *mroQ* mutant strain (**Fig 5**). Although an obvious explanation for this phenotype is not immediately clear, in addition to regulation of virulence factors, AgrA has previously been shown to have a role in regulating certain metabolic genes (11), some of which are indicated to have a role in staphyloxanthin production (11, 36).

**Figure 9.**
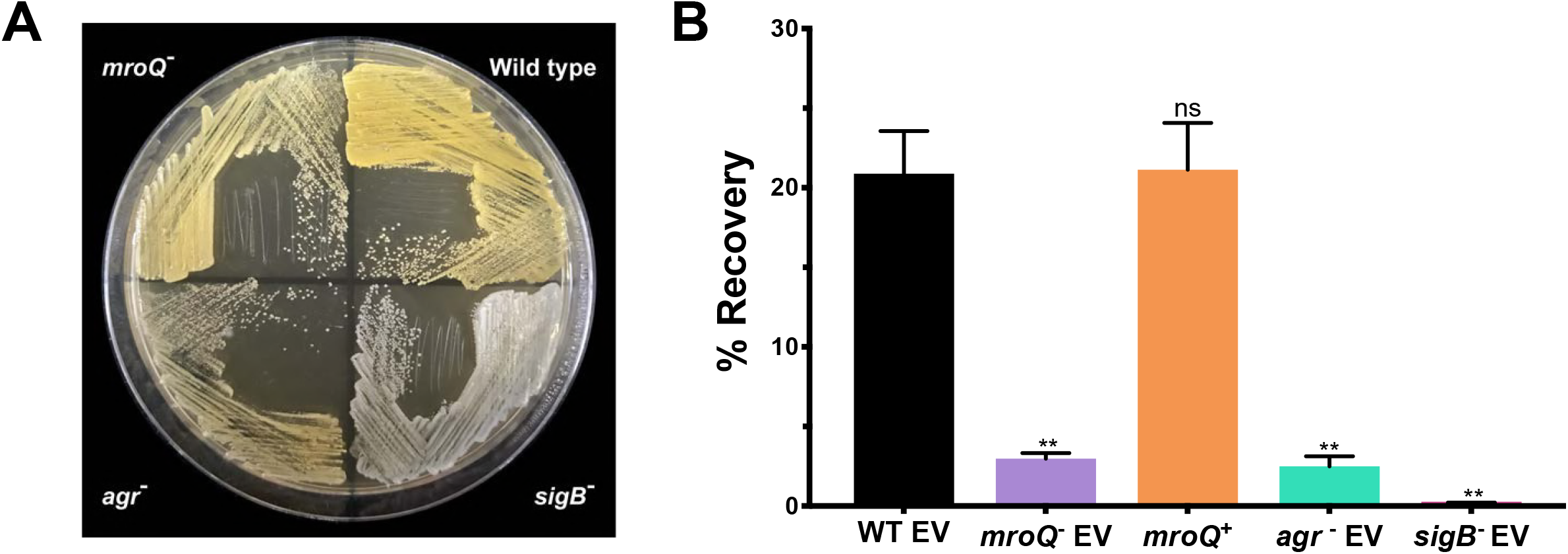
Staphyloxanthin Production and Tolerance to Oxidative Stress is Decreased in the *mroQ* Mutant in an *agr* Dependent Manner. The strains indicated were streaked onto TSA and grown overnight (**A**). Percent recovery of strains following 30 min exposure to H_2_O_2_ at a 20 mM concentration (**B**). A Student’s t-test was used to determine statistical significance, ** = p ≤ 0.01, n.s. = not significant. Error bars shown with SEM. EV = Empty Vector.

Due to the importance of staphyloxanthin in protecting *S. aureus* against reactive oxygen species, we performed oxidative stress assays with our collection of strains (**Fig 9B**) alongside a *sigB* null mutant as a pigment-lacking control. Accordingly, strains were exposed to hydrogen peroxide (H_2_O_2_) at a concentration of 20 mM for 30 minutes. Following this, CFU/ml was determined and compared to the inocula, producing percent recovery for each strain. As expected, the *sigB* knockout strain demonstrated only a 0.18% survival under these conditions, whilst the wild type strain produced a 20.9% recovery. The *mroQ* mutant demonstrated 3.0% recovery in comparison to our parental strain, whilst the *agr* mutant demonstrated 2.5% recovery. Importantly, complementation of the *mroQ* mutant resulted in wild type levels of survival under these conditions. These findings indicate that, in accordance with decreased staphyloxanthin production, the *mroQ* and *agr* mutants both demonstrate reduced tolerance to oxidative stress.

### MroQ elicits protease dependent control of Agr activity

As previously discussed, Abi-domain proteins also belong to the M79 metalloprotease family (25, 27, 37). These enzymes are defined by several conserved motifs, with the primary being a catalytic dyad comprised of neighboring glutamic acid residues (**Supplemental Figure S1**) (25). Thus far, the conserved protease residues of Abi-domain proteins have not been demonstrated as having essential roles in their function in previous studies involving Gram-positive bacteria (27, 30). As such, we sought to investigate the importance of proteolytic activity for MroQ in exerting its Agr modulating effects. To do this we created a catalytically inert complementing strain, where the proteolytic residues were converted from glutamic acid to alanine (E141A/E142A) (**Supplemental Figure S3A**). Upon introducing this construct into the *mroQ* mutant strain we found that this site-directed-mutant complement did not restore proteolytic or hemolytic activity to *S. aureus* (**Fig 1A, Fig 2A**), demonstrating levels equivalent to the *mroQ* and *agr* mutant strains. As such, these results indicate that MroQ is the first Gram-positive Abi protein to be dependent on its proteolytic capacity for function.

### The presence of mature AIP in *mroQ* mutant supernatants indicates it plays no role in the processing of AgrD

Given that MroQ controls *agr* activity in a protease dependent manner, and that a central component of the Agr system (the AgrD pheromone-precursor) is subject to proteolytic processing, we set out to explore whether MroQ is required for AIP maturation. As such, we used UPLC-MS/MS analysis to identify mature AIP in the culture supernatants of our various strains. In so doing, we observed AIP-I (Retention Time = 2.20-2.23 minutes) in culture supernatants for the wild type, *mroQ* mutant, and *mroQ* complemented strains, but not in the *agr* mutant or in the TSB blank control (**Fig 10A**). Importantly, when analyzing fragmentation patterns, the expected fragments of AIP-I (38) were also observed in supernatants of the wild-type and *mroQ* mutant (**Fig 10B**), but not for the agr-null strain. Next, the area under the curve (AA) of AIP peaks in Figure 10A, in conjunction with an AIP-I standard curve, was used to calculate AIP-I concentrations in the various culture supernatants (38) (**Supplemental Table S5**). Although the *mroQ* mutant was below the lower threshold for accurate quantification (0.1 ng/ml), extrapolation analysis suggests that secretion of mature AIP-I is ~50 fold lower in the mutant relative to the wild type (AA = 260301 and 3224, for wild type and *mroQ* mutant, respectively).

**Figure 10.**
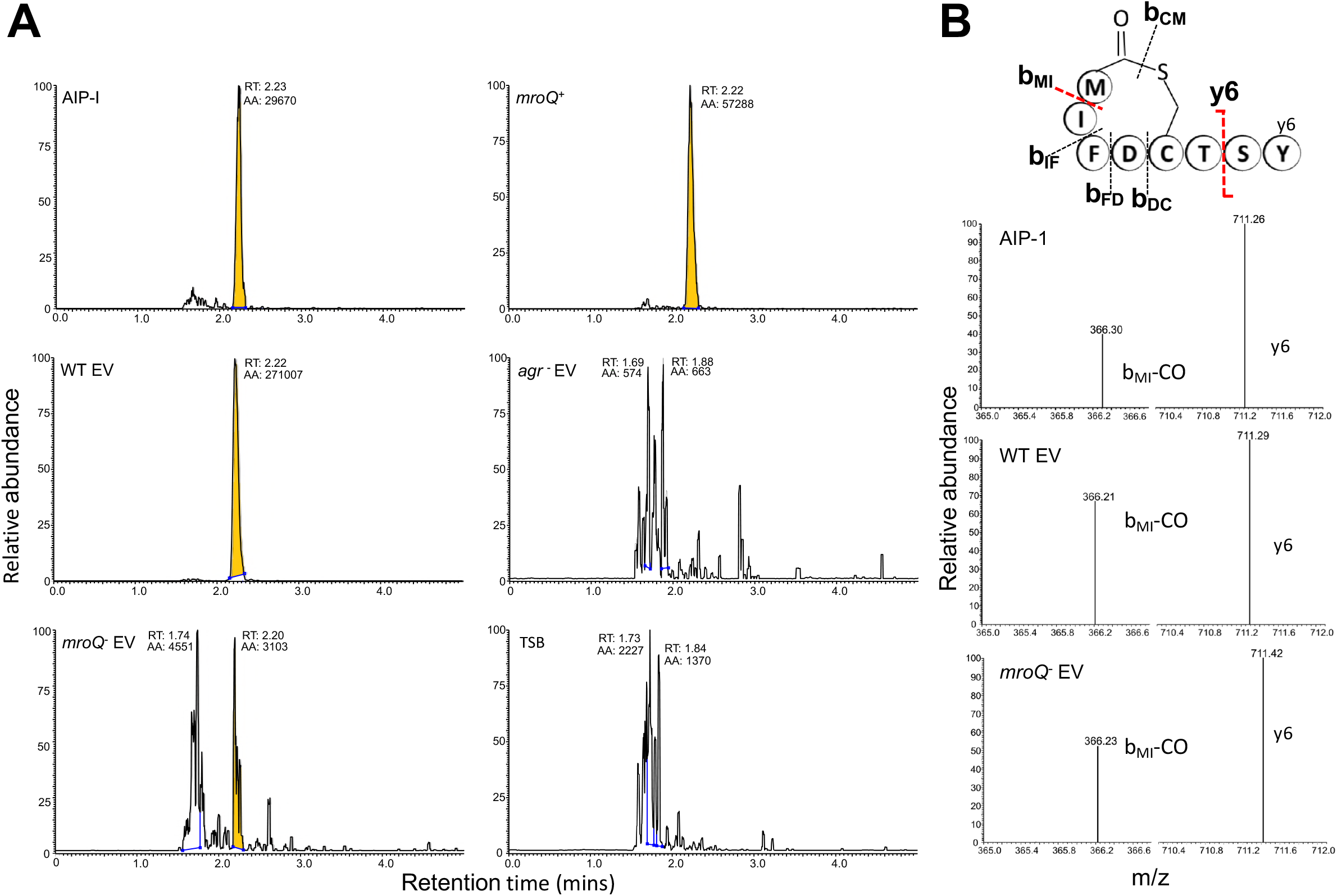
Despite Impaired *agr* Activity the *mroQ* Mutant Still Produces Fully Mature AIP. Culture supernatants from the wild type, *mroQ^−^* mutant, *agr* mutant, and the *mroQ^+^* complement strains, alongside sterile TSB with or without purified AIP-I, underwent UPLC-MS/MS analysis. Experiments were performed three times for each condition, with chromatograms and spectra shown herein proving representative. Chromatogram showing elution of peptides matching the AIP standard are presented. Peaks highlighted in yellow at retention time (RT) 2.20-2.23 minutes correspond to mature AIP-I, and for all conditions peaks with ~100 % relative abundance are labelled with RT and the peak area (AA) (**A**). MS/MS spectra comparing the relative abundance of the MRM transitions (*m/z* 961.3 to 711.2 and 365.9) at the retention time of ~2.20 for 0.1 ng/ml AIP-I standard, wild type (WT) and *mroQ^−^* mutant, shown below a schematic of AIP-I with the transitions indicated by dotted red lines (**B**). The first enables liberation of the y6 ion of AIP (calculated *m/z* 711.30), whilst the second fragments the cyclic part of AIP, bMI, a prevalent fragment of the cyclic component of AIP (calculated *m/z* 366.23). EV = Empty Vector.

As such, it appears that MroQ does not exert its protease-dependent effects on *agr* activity via direct processing of AgrD, or by preventing the proteolytic activity of AgrB (or SpsB, which is also required for AgrD cleavage (39, 40)).

### MroQ plays an important role during abscess formation in both localized and systemic infections

Using murine models of infection, we compared the virulent capacity of the *mroQ* and *agr* mutants alongside our wild type strain in systemic and localized infections. Upon assessing the mortality of mice infected with each strain during a murine model of sepsis and dissemination, we observed no overall difference in survival across the three strains (data not shown). This is perhaps not surprising given that the *agr* system has previously been shown to have a limited role in influencing this kind of infectious capacity (4, 41). Interestingly, however, we noted that the kidneys of mice infected with both mutant strains, whilst having no alteration in bacterial burden (**Fig 11A**), had an increase in abscess formation (**Fig 11B**). Although curious, it has previously been shown that a defective Agr system results in the increased production of surface factors that are important to abscess formation, whilst at the same time limiting production of secreted factors required for dissemination from the abscess (42). As such, this would explain why the *mroQ* mutant demonstrates an increase in renal abscess formation compared to wild type, which mirrors, albeit to a lesser extent, that of a complete *agr* deletion strain.

**Figure 11.**
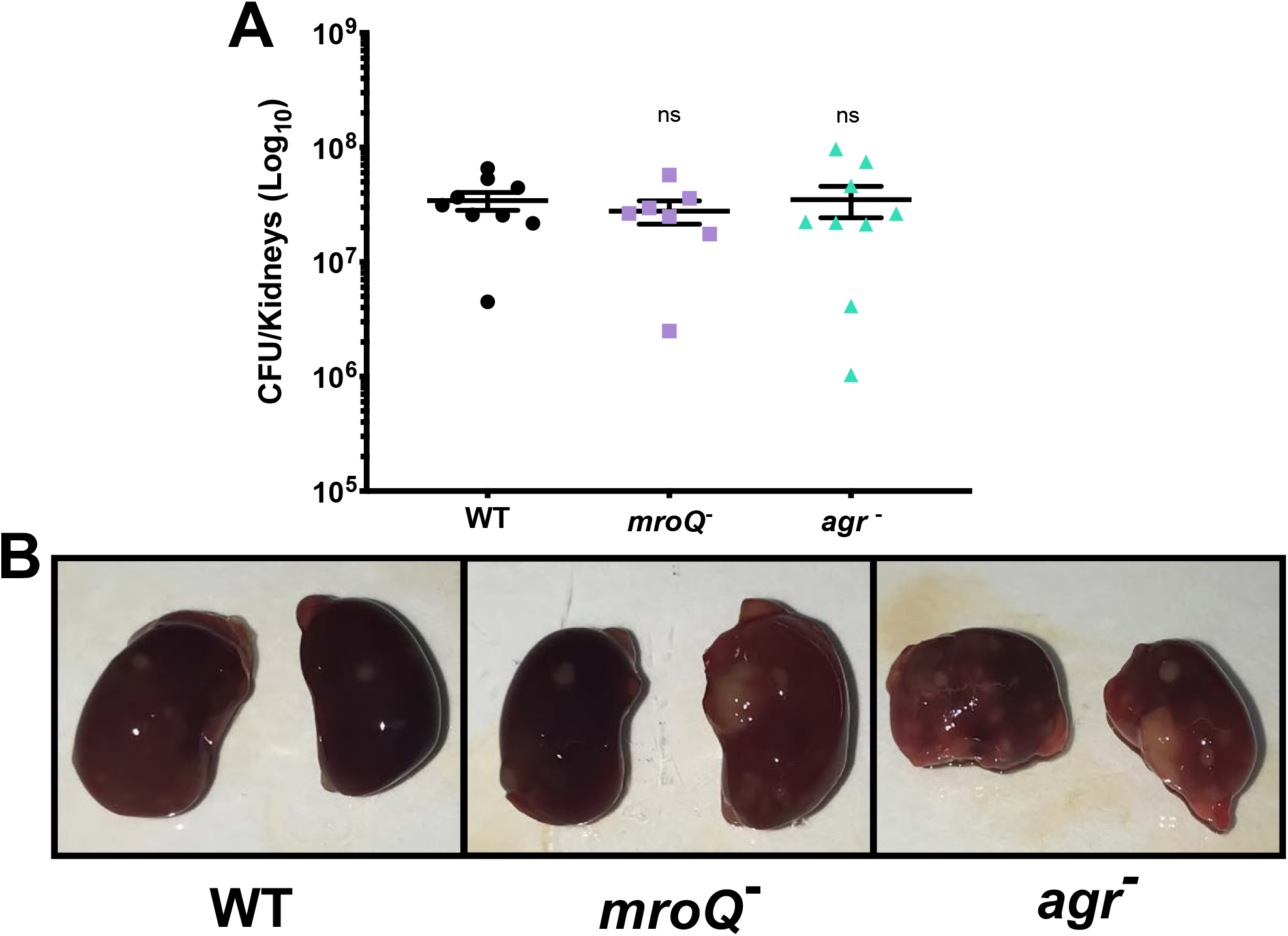
Mutation of *mroQ* Results in Increased Renal Abscess Formation. Female, 6-week old CD-1 mice were injected with the strains shown, and infections were allowed to proceed for 7 days. Following the infection period, mice were sacrificed, and the kidneys were harvested. Organs were homogenized in 1 ml of PBS, serially diluted, and plated on TSA to determine bacterial burden (**A**). A Mann-Whitney test was used to determine statistical significance, n.s. = not significant. Representative images of kidneys harvested from mice infected with the strains noted (**B**).

Skin and soft tissue infections (SSTIs) are commonly caused by *S. aureus*, particularly by USA300 isolates, which are the predominant cause of SSTIs in the United States (43). The success of CA-MRSA strains in causing SSTIs is a result of their overproduction of important virulence factors (Hla, PSMs, secreted proteases), arising from a hyperactive Agr system (44). To investigate the requirement for MroQ in this kind of localized infections, we next used a murine model of subcutaneous abscess formation with our collection of strains. Following the 7 day infection period, mice were sacrificed, abscesses were excised and measured, and bacterial burden determined for each abscess. Upon measuring abscess area (mm^2^), the *mroQ* mutant had a 2-fold decrease in comparison to wild type, whilst the *agr* mutant had a 4.5-fold decrease (**Fig 12A**). Additionally, the *mroQ* mutant demonstrated decreased bacterial recovery in the abscess model (5.2-fold decrease) compared to the parent, which, although significant, was not at the same magnitude as the *agr* mutant (212.2-fold decrease) (**Fig 12B**). Finally, the *mroQ* mutant presented decreased levels of dermonecrosis compared to the parental strain, whilst the *agr* mutant produced no detectable areas of dermonecrosis at all (**Supplemental Figure S4**). This finding is perhaps to be expected given that this kind of pathology has previously been shown to heavily depend on the Agr regulated factors (43, 45). Collectively, *mroQ* disruption results in attenuated virulence and dermonecrosis in models of localized skin infection that likely arises from impaired, although not ablated, Agr activity within this strain.

**Figure 12.**
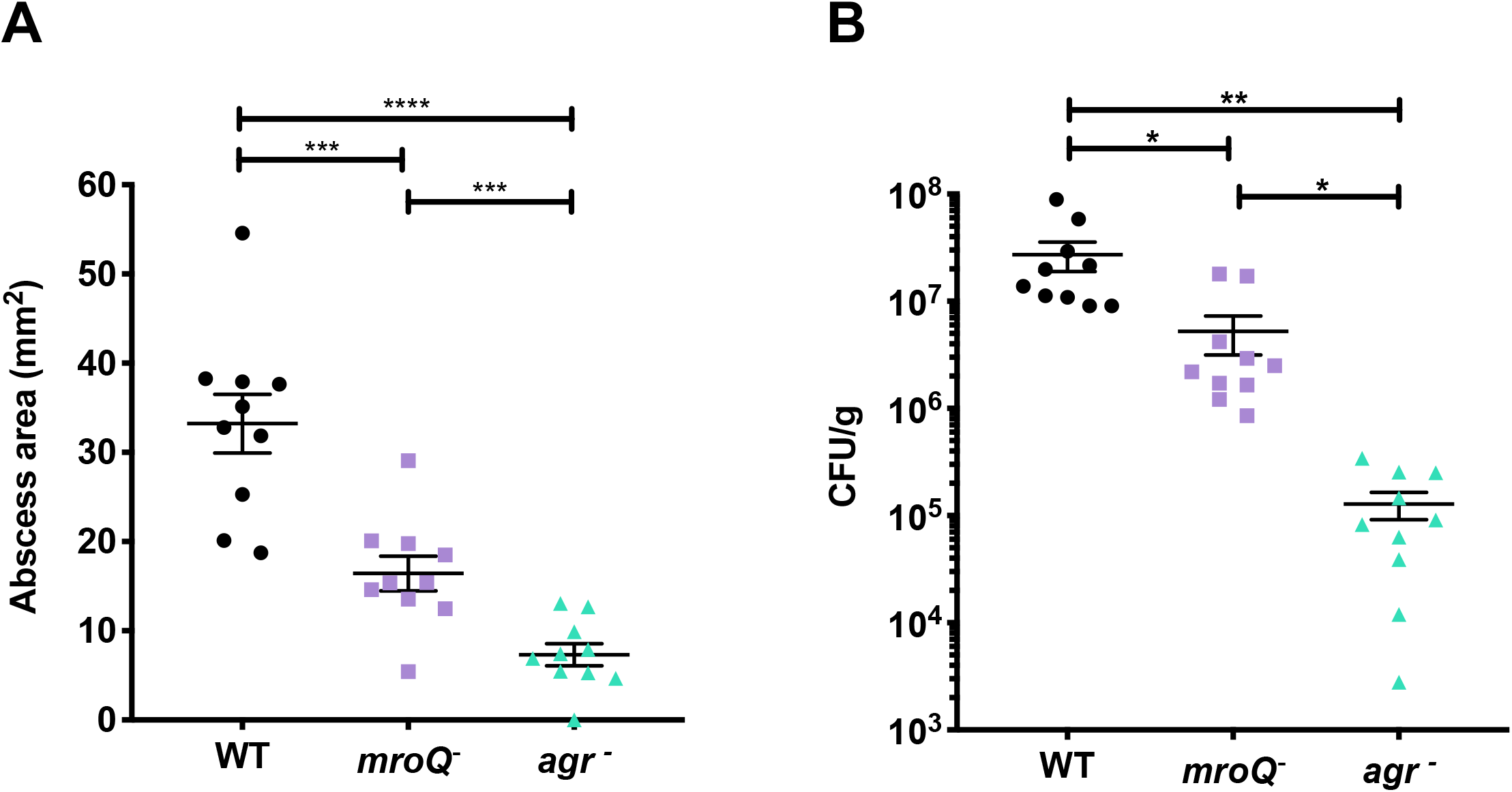
The *mroQ* Mutant Demonstrates Attenuated Virulence in a Murine Abscess Model of Skin Infection. Female, 6-week old BACL/c mice were injected subcutaneously, and infections were allowed to proceed for 7 days. Abscesses were measured (width and height), and area was calculated (mm^2^). (**A**). Bacterial burden was determined for each abscess (CFU/g) (**B**). A Student’s t-test was used to determine statistical significance, *= p<0.05, ** = p ≤ 0.01, *** = p ≤ 0.001, **** = p ≤ 0.0001. Error bars shown with SEM.

## Discussion

Numerous factors have to date been identified as playing a role in the regulation of Agr activity, including transcription factors, antisense RNAs, and host factors (5, 8, 20, 22, 23, 46). In this study, we characterize a novel Abi-domain protein, and its previously unidentified role as a novel regulator of Agr. Whilst exploring the impact of MroQ on *agr* function, we examined the expression of numerous known regulators of the *agr* system (**Fig 13**), including activators and repressors (5, 20, 47). In sum, only one substantial alteration was observed amongst the collected repressors, which was for *sarR* transcription, although this change was in fact a decrease rather than upregulation. Regarding activators, some Sar-family member protein, including MgrA and SarA, did demonstrate decreased expression, however these effects were only ~2-3-fold in magnitude. As such, the lack of significant changes observed in key repressors and activators does not appear to explain the substantial decreases observed in Agr activity upon *mroQ* disruption. These findings suggest that the influence of MroQ on Agr activity is likely to be occurring through direct interaction with Agr proteins, or through as yet unknown effectors of the system.

**Figure 13.**
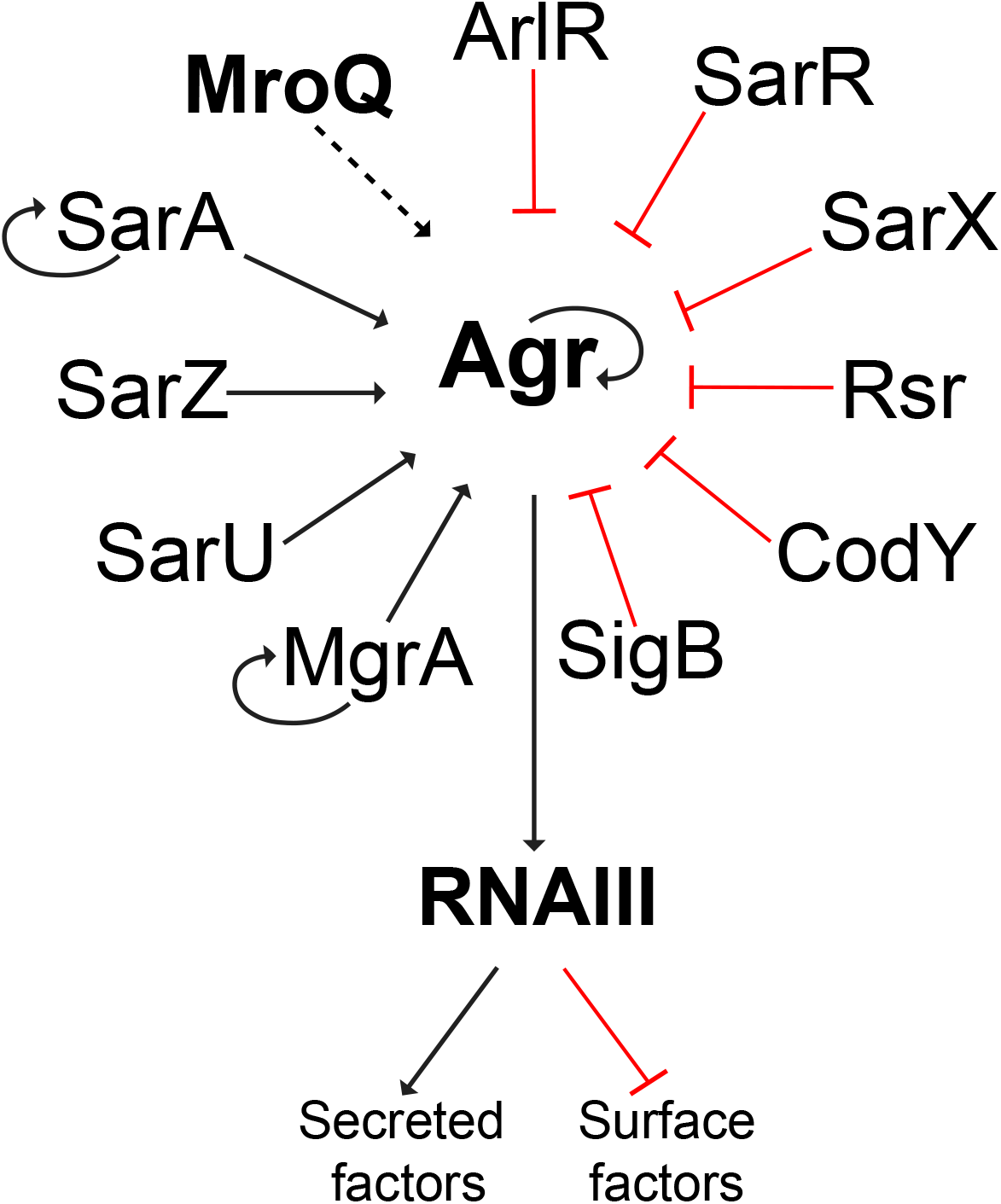
Regulatory Map Illustrating Known Direct Regulators of Agr Activity. A map of *agr* regulators was developed based on the literature. Changes in expression of these regulators does not correlate with the substantial decrease in Agr activity observed upon *mroQ* disruption. MroQ appears to directly interact with Agr, resulting in a downstream effect on virulence factor production. Black arrows indicate activation, red crossbar indicates repression, and black dotted arrow indicates putative activation.

How such a direct effect could be mediated is perhaps suggested by existing literature on other bacterial Abi proteins. In the first of two recent accounts characterizing Abi-domain proteins in Gram-positive bacteria, a protein called Abx1 was found to have a role in virulence gene expression in *Streptococcus agalactiae* (27). Using a two-hybrid study, it was revealed that Abx1 interacts with the CovS histidine kinase, seemingly via the transmembrane domains of this protein (27). Of note, this interaction was not dependent on the proteolytic activity of Abx1, as mutation of its conserved Glu-Glu residues did not affect function towards the CovRS system. This study represented a first of its kind, elucidating a novel function for Abi-domain proteins in Gram-positive bacteria. More recently, another Abi-domain protein was characterized in *S. aureus* (30). Here, Poupel *et al* investigated SpdC, a membrane protein involved in an interaction network controlling the coordination of division with cell envelope metabolism and host interactions (30). In this study, it was determined that SpdC interacts with 10 different histidine kinases, with several interactions being through their transmembrane domains, similar to the findings in GBS (27). Importantly, SpdC is one of the three Abi proteins in *S. aureus* (**Supplemental Figure S1**) to lack the conserved EE catalytic residues, once again showing that proteolytic activity is not essential for its HK regulating role.

Thus far, our findings suggest that MroQ may interact directly with the Agr system to modulate its activity. The *mroQ* gene is encoded ~4 kb upstream of the *agr* operon and is expected (as with other Abi proteins) to specify a membrane protein. Importantly, when we performed localization studies for MroQ we did indeed find that it is only present in membrane fractions (**Supplemental Figure S3B**). At this time, we have yet to confirm potential partners of interaction for MroQ, however, considering the growing evidence demonstrating Abi-domain proteins in Gram-positive pathogens interact with and regulate histidine kinases (27, 30), we suggest that MroQ could perhaps mediate its function through AgrC. We note that the Agr system also encodes another membrane protein, AgrB, and that this protein itself also functions as a protease (processing the AgrD pheromone precursor). Our data at this time does not, however, implicate AgrB as the target of MroQ. Specifically, our analysis of culture supernatants by mass spectrometry detected the presence of mature AIP as being produced by the *mroQ* mutant. As such, this would suggest that MroQ has no role in directly processing AgrD and is not required for AgrB to function correctly. Similarly, we believe it is unlikely that MroQ is involved in processing AgrA as AgrC-AgrA interactions have been extensively studied through various crystallization, biochemical, and biophysical studies and do not appear to need adapter proteins to engage (48).

How MroQ might interact with its target(s) within the *S. aureus* cell is currently unclear, but it is seemingly dependent on its proteolytic activity. The M79 peptidase family in eukaryotes typically cleave C-terminal tripeptides from isoprenylated proteins (49), with known substrates including Ras and Ras-related GTP-binding proteins, and protein kinases (50). This latter category again provides a link between Abi/M79 proteins and kinase enzymes, further suggesting that AgrC may indeed be the target of MroQ. Determining if the C-terminus of AgrC is subject to proteolysis for correct function is an important consideration for continued exploration of MroQ and its role within the *S. aureus* cell.

Using RNA-seq analysis, we found that the entire *agr* system was downregulated in our *mroQ* mutant, with the most substantial decreases observed for *agrC* and *agrA*, the two-component system. When comparing this dataset to a similar transcriptome generated with an *agr* operon deletion mutant we observed 55% similarity in transcriptome between the two strains, with 25% of these changes corresponding to virulence factors. Of note, a large number of these virulence factors, although substantially altered in the *mroQ* mutant, were changed to a much greater degree in the *agr* mutant (**Table 3**). This is seen on a greater scale when observing trends in virulence factor production where proteolytic and hemolytic activity were decreased but not completely diminished in the *mroQ* null strain, as well as in virulence with our murine model of skin infection. Given that *mroQ* mutation phenocopies, but does not completely replicate, the act of *agr* deletion, we suggest that this protein is necessary, but not sufficient, for Agr activity in *S. aureus*.

A singular concern presents itself in the context of our suggestion that mutation of *mroQ* creates a diminished but not abrogated *agr* phenotype: That our phenotype might arise from the fact that we are using a mutational insertion strain, rather than a deletion mutant. We are supported by the observation that, although there have been screens performed to uncover *S. aureus* mutants that result in ahemolytic phenotypes (51), MroQ has not been identified to date. However, to explore this more fully we reviewed our *mroQ* mutant RNA-sequencing data set and determined that transposon insertion successfully terminates the *mroQ* transcript at the point of insertion (**Supplemental Figure S5**).

In the context of our virulence studies, we made a striking observation when considering the gross pathology presented in the kidneys of mice using a sepsis model. Here, abscess frequency and severity were increased in the *mroQ* mutant, and to an even greater level in the *agr* null strain. At first this might seem contradictory, however when one explores the literature regarding *S. aureus* abscess formation, this becomes more logical. Specifically, it is known that the early phases of abscess formation are largely dependent on Agr repressed surface factors, such as Spa, SdrD, ClfA, and Coa. Furthermore, the later stages of infection are characterized by the rupturing of abscess to elicit dissemination, a process known to be dependent on Agr-activated factors such as secreted toxins and proteases (2, 52, 53). Accordingly, a defect in quorum sensing, as observed in the *mroQ* mutant, locks abscess formation in the early stages, preventing development to later phases, which are characterized by necrosis and dissemination to new sites (45). The finding that this is more severe in the *agr* mutant than the *mroQ* null strain is again in line with our notion that MroQ is a factor that contributes to Agr activity, but is not the only factor so doing.

As previously mentioned, models for skin infection are often used to represent SSTIs, which are commonly caused by USA300 strains (54). A key hallmark of subcutaneous skin infections by such strains is dermonecrosis, which presents as necrosis of the epidermis and dermis, and has previously been shown to heavily depend on the Agr regulated factors α-hemolysin (Hla) and phenol soluble modulins (PSMs) (43, 45, 55). Importantly, our *mroQ* mutant demonstrated decreased promoter activity for *hla* (3.7-fold decrease), although not to the level of an *agr* mutant (6.4-fold decrease). These transcription-based findings directly translate into virulence-based outcomes, as dermonecrosis presented by the *mroQ* mutant, although reduced compared to the wild type, was not absent, as seen for the *agr* mutant. Similarly reduced-but-not-ablated findings for the *mroQ* mutant were presented from our study of abscess size and bacterial load per abscess, where again the *agr* mutant was profoundly impaired in its pathogenic potential compared to the parent.

Agr-related factors also play a largely identical role in biofilm formation as they do in abscess formation, which explains why *agr* defective strains demonstrate increased formation of biofilms (53). Biofilms undergo a cycle comprised of three stages: initial attachment, maturation, and dispersal (53, 56). During the initial attachment, adhesin proteins play a crucial role in attachment to abiotic or biotic surface, as well as in the maturation phase, where they are important for cell-cell adhesion. Conversely, the dispersal of biofilms is highly dependent on enzymes such as proteases and nucleases produced by the bacteria to break down components of the biofilm matrix, ultimately allowing for the continuation of the biofilm life cycle from planktonic cells (53). Our *mroQ* mutant demonstrates excellent initial attachment and maturation, likely due to its increased production of MSCRAMMs, however fails to properly disperse due to decreased Agr activity, and therefore decreased production of secreted enzymes necessary for breaking down the extracellular matrix (53).

In summary, this study identifies MroQ, a novel transmembrane protein, as having a critical role in regulation of virulence factor production via modulation of the *agr* quorum sensing system. Further studies are required in order to elucidate the mechanism by which modulation occurs, and more specifically to identify MroQ partner(s) of interaction. Moreover, we must determine how proteolysis is necessary to the function of MroQ, especially considering this is the first characterized Abi-domain protein to demonstrate protease dependent activity. In addition, four other Abi-domain proteins exist in *S. aureus*, SAUSA300_0932, SAUSA300_2405, SpdA, and SpdB, the latter two of which have been shown to have a role in surface protein regulation (37). Given that surface protein display is under the control of several TCS in *S. aureus* (57–59) it is entirely possible that they may function by interaction with HKs as well. Accordingly, the continued characterization of MroQ, and the complete set of Abi-domain proteins in *S. aureus*, remains a focus of study in our laboratory.

## Materials and Methods

### Reagents, media, and growth conditions

All bacterial strains and plasmids utilized in this study are listed in Table 1. Bacterial strains were grown at 37°C with shaking at 250 rpm. *S. aureus* bacterial strains were grown in tryptic soy broth (TSB) or on tryptic soy agar (TSA), whilst *E. coli* was grown in lysogeny broth (LB) or on LB agar. TSB supplemented with 5% human plasma was utilized for biofilm experiments. Some investigations required the use of antibiotics for selection, which were added at the following concentrations for *S. aureus:* Erythromycin and tetracycline, 5 μg/ml, lincomycin, 25 μg/ml, chloramphenicol, 10 μg/ml. For *E. coli* strains, ampicillin was used at a concentration of 100 μg/ml. When necessary, X-gal was utilized at a concentration of 40 μg/ml. The induction of complementation was achieved using 0.4 μM cadmium chloride (CdCl_2_) to induce protein expression. Throughout this study, strains were grown overnight, followed by a 1:100 dilution into fresh TSB and additional growth for 3 h. Strains were then standardized to the OD_600_ required for each experiment (detailed below).

**Table 1.**
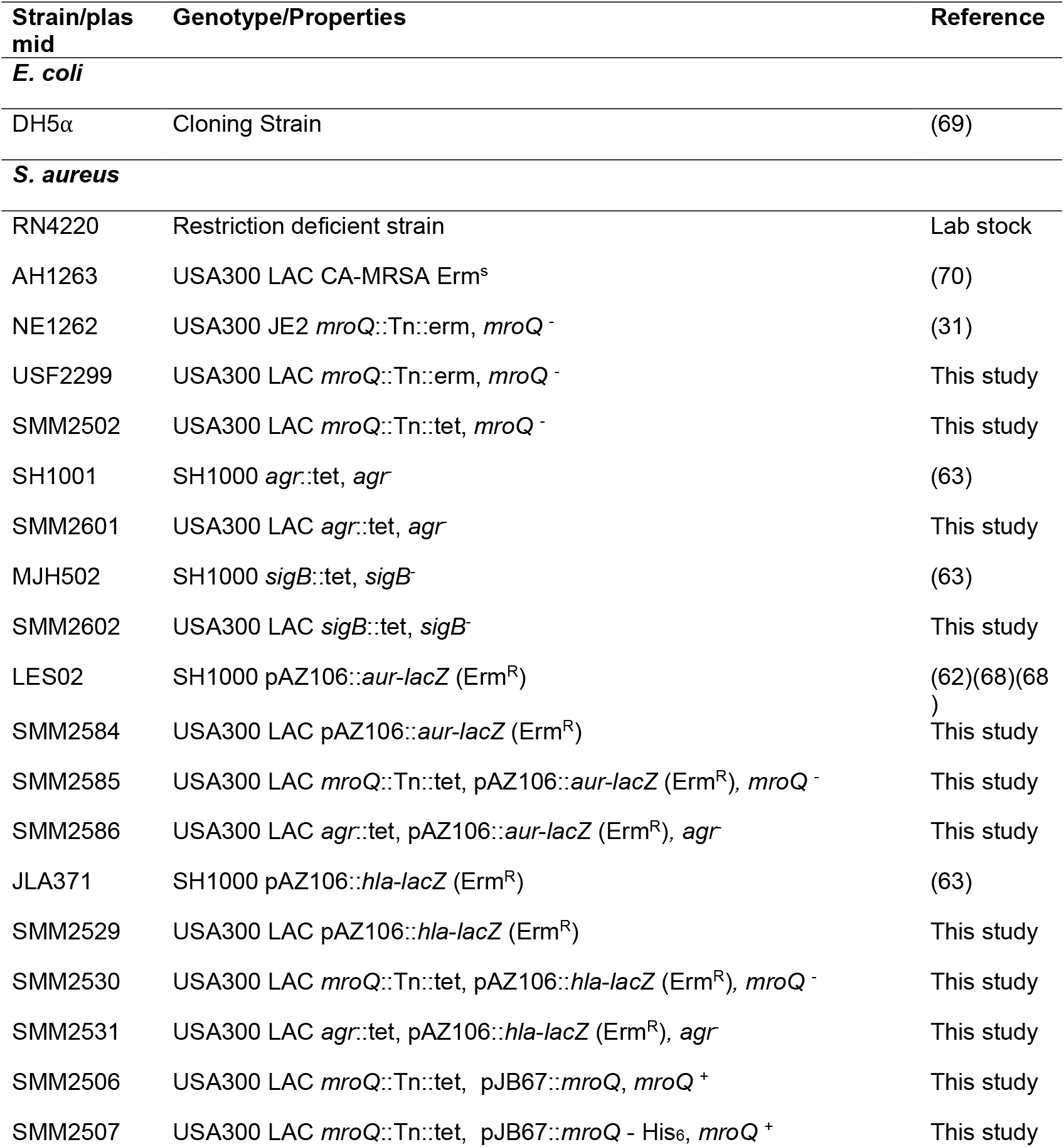
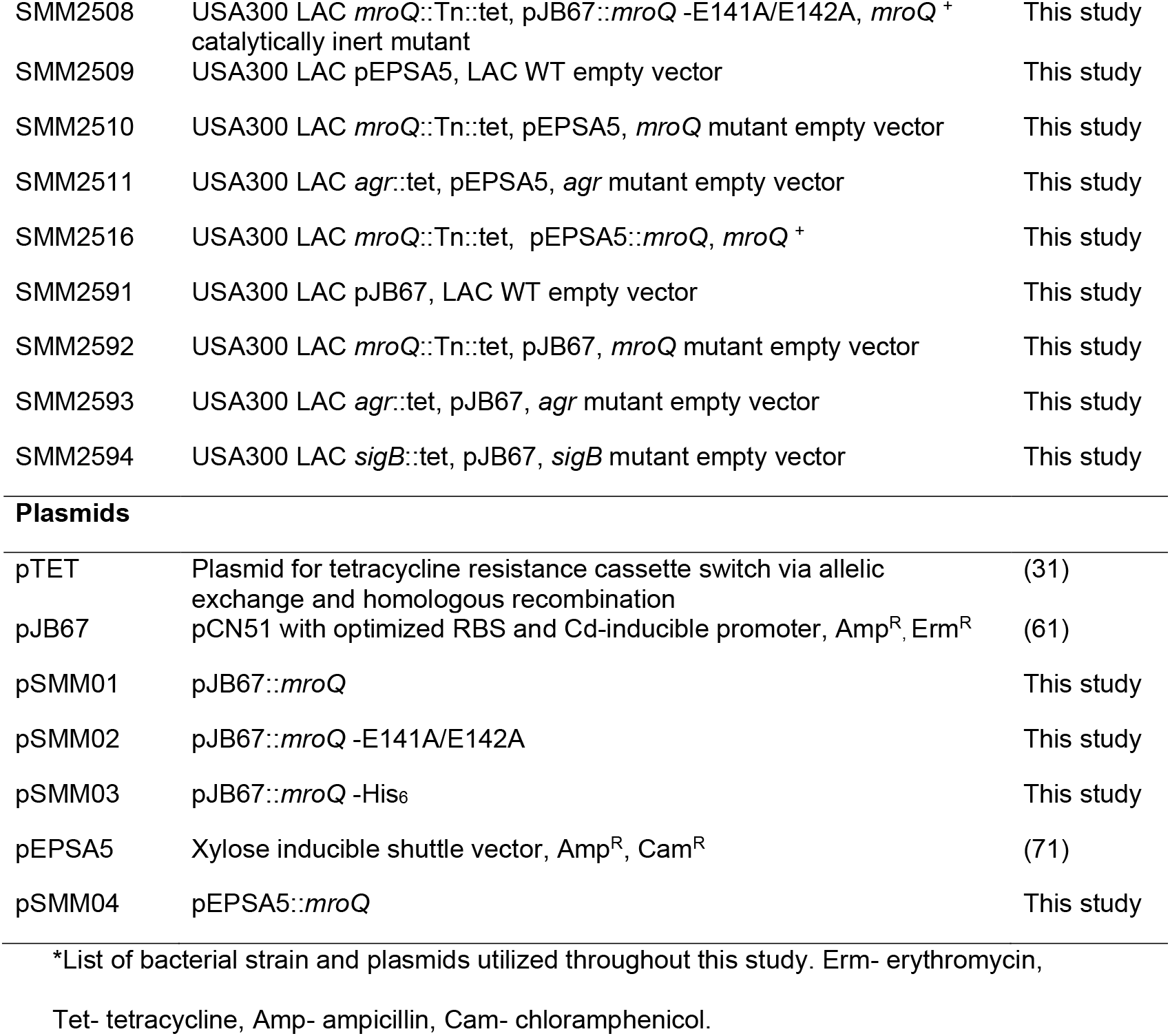
Strains and plasmids used in this study.

| Strain/plasmid | Genotype/Properties | Reference |
| --- | --- | --- |
| ***E. coli*** |  |  |
| DH5α | Cloning Strain | (69) |
| ***S. aureus*** |  |  |
| RN4220 | Restriction deficient strain | Lab stock |
| AH1263 | USA300 LAC CA-MRSA Erm<sup>s</sup> | (70) |
| NE1262 | USA300 JE2 *mroQ*::Tn::erm, *mroQ*<sup>-</sup> | (31) |
| USF2299 | USA300 LAC *mroQ*::Tn::erm, *mroQ*<sup>-</sup> | This study |
| SMM2502 | USA300 LAC *mroQ*::Tn::tet, *mroQ*<sup>-</sup> | This study |
| SH1001 | SH1000 *agr*::tet, *agr* | (63) |
| SMM2601 | USA300 LAC *agr*::tet, *agr* | This study |
| MJH502 | SH1000 *sigB*::tet, *sigB* | (63) |
| SMM2602 | USA300 LAC *sigB*::tet, *sigB* | This study |
| LES02 | SH1000 pAZ106::*aur-lacZ* (Erm<sup>R</sup>) | (62)(68)(68) |
| SMM2584 | USA300 LAC pAZ106::*aur-lacZ* (Erm<sup>R</sup>) | This study |
| SMM2585 | USA300 LAC *mroQ*::Tn::tet, pAZ106::*aur-lacZ* (Erm<sup>R</sup>), *mroQ*<sup>-</sup> | This study |
| SMM2586 | USA300 LAC *agr*::tet, pAZ106::*aur-lacZ* (Erm<sup>R</sup>), *agr* | This study |
| JLA371 | SH1000 pAZ106::*hla-lacZ* (Erm<sup>R</sup>) | (63) |
| SMM2529 | USA300 LAC pAZ106::*hla-lacZ* (Erm<sup>R</sup>) | This study |
| SMM2530 | USA300 LAC *mroQ*::Tn::tet, pAZ106::*hla-lacZ* (Erm<sup>R</sup>), *mroQ*<sup>-</sup> | This study |
| SMM2531 | USA300 LAC *agr*::tet, pAZ106::*hla-lacZ* (Erm<sup>R</sup>), *agr* | This study |
| SMM2506 | USA300 LAC *mroQ*::Tn::tet, pJB67::*mroQ*, *mroQ*<sup>+</sup> | This study |
| SMM2507 | USA300 LAC *mroQ*::Tn::tet, pJB67::*mroQ* - His<sub>6</sub>, *mroQ*<sup>+</sup> | This study |
| SMM2508 | USA300 LAC *mroQ*::Tn::tet, pJB67::*mroQ* -E141A/E142A, *mroQ* <sup>+</sup> catalytically inert mutant | This study |
| SMM2509 | USA300 LAC pEPSA5, LAC WT empty vector | This study |
| SMM2510 | USA300 LAC *mroQ*::Tn::tet, pEPSA5, *mroQ* mutant empty vector | This study |
| SMM2511 | USA300 LAC *agr*::tet, pEPSA5, *agr* mutant empty vector | This study |
| SMM2516 | USA300 LAC *mroQ*::Tn::tet, pEPSA5::*mroQ*, *mroQ* <sup>+</sup> | This study |
| SMM2591 | USA300 LAC pJB67, LAC WT empty vector | This study |
| SMM2592 | USA300 LAC *mroQ*::Tn::tet, pJB67, *mroQ* mutant empty vector | This study |
| SMM2593 | USA300 LAC *agr*::tet, pJB67, *agr* mutant empty vector | This study |
| SMM2594 | USA300 LAC *sigB*::tet, pJB67, *sigB* mutant empty vector | This study |
| Plasmids |  |  |
| pTET | Plasmid for tetracycline resistance cassette switch via allelic exchange and homologous recombination | (31) |
| pJB67 | pCN51 with optimized RBS and Cd-inducible promoter, Amp<sup>R</sup>, Erm<sup>R</sup> | (61) |
| pSMM01 | pJB67::*mroQ* | This study |
| pSMM02 | pJB67::*mroQ* -E141A/E142A | This study |
| pSMM03 | pJB67::*mroQ* -His<sub>6</sub> | This study |
| pEPSA5 | Xylose inducible shuttle vector, Amp<sup>R</sup>, Cam<sup>R</sup> | (71) |
| pSMM04 | pEPSA5::*mroQ* | This study |

\*List of bacterial strain and plasmids utilized throughout this study. Erm- erythromycin,
Tet- tetracycline, Amp- ampicillin, Cam- chloramphenicol.

### Mutant strain construction

A *mroQ* mutant was obtained from the Nebraska transposon mutant library (NTML) (31). The USA300 JE2 SAUSA300_1984 transposon mutant was used to create a phage lysate using phi11 bacteriophage (Φ11), which was subsequently transduced into the *S. aureus* USA300 LAC background and confirmed by PCR using gene specific primers (Table 2). Antibiotic resistance cassette switching for the *mroQ* mutant, from erythromycin to tetracycline, was achieved via allelic exchange and homologous recombination as described by Bose and coworkers (31). These manipulations were again confirmed via PCR, followed by transduction into a clean USA300 LAC background (and a subsequent round of PCR confirmation). Additionally, SH1000 *agr* and SH1000 *sigB* mutants were used to create phage lysates for transduction into the *S. aureus* USA300 LAC background.

**Table 2.**
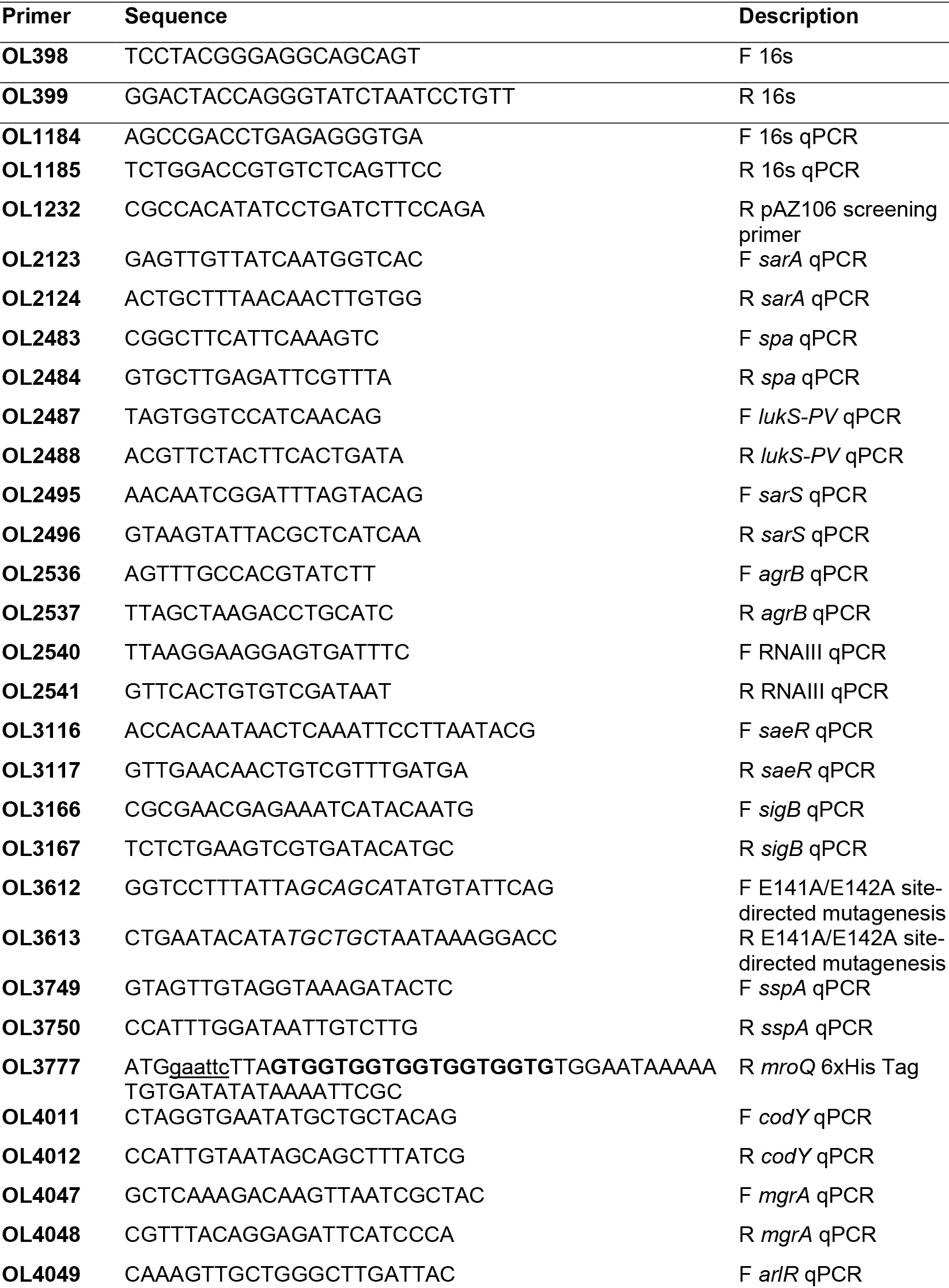
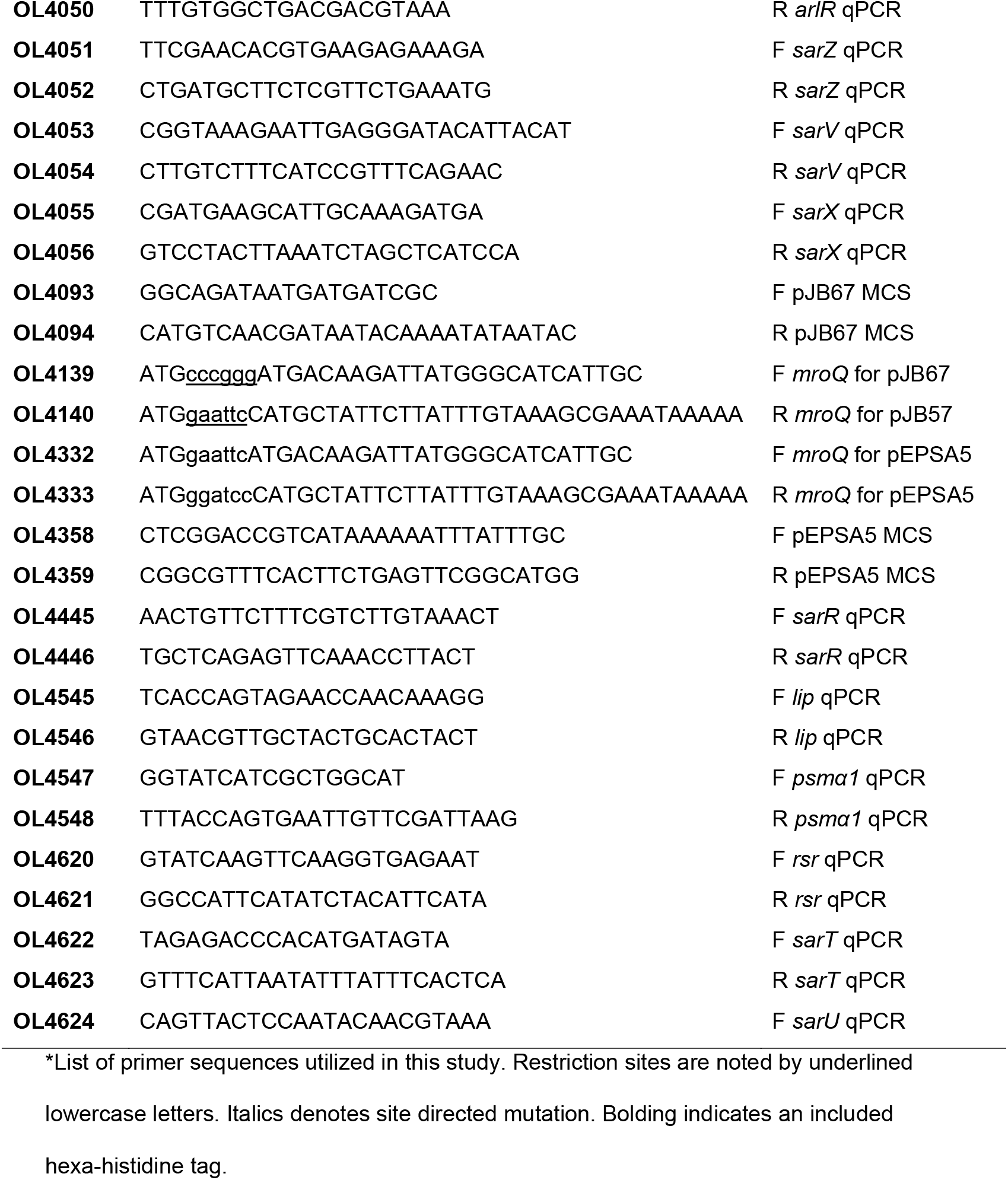
Primers used in this study.

**Table 3.**
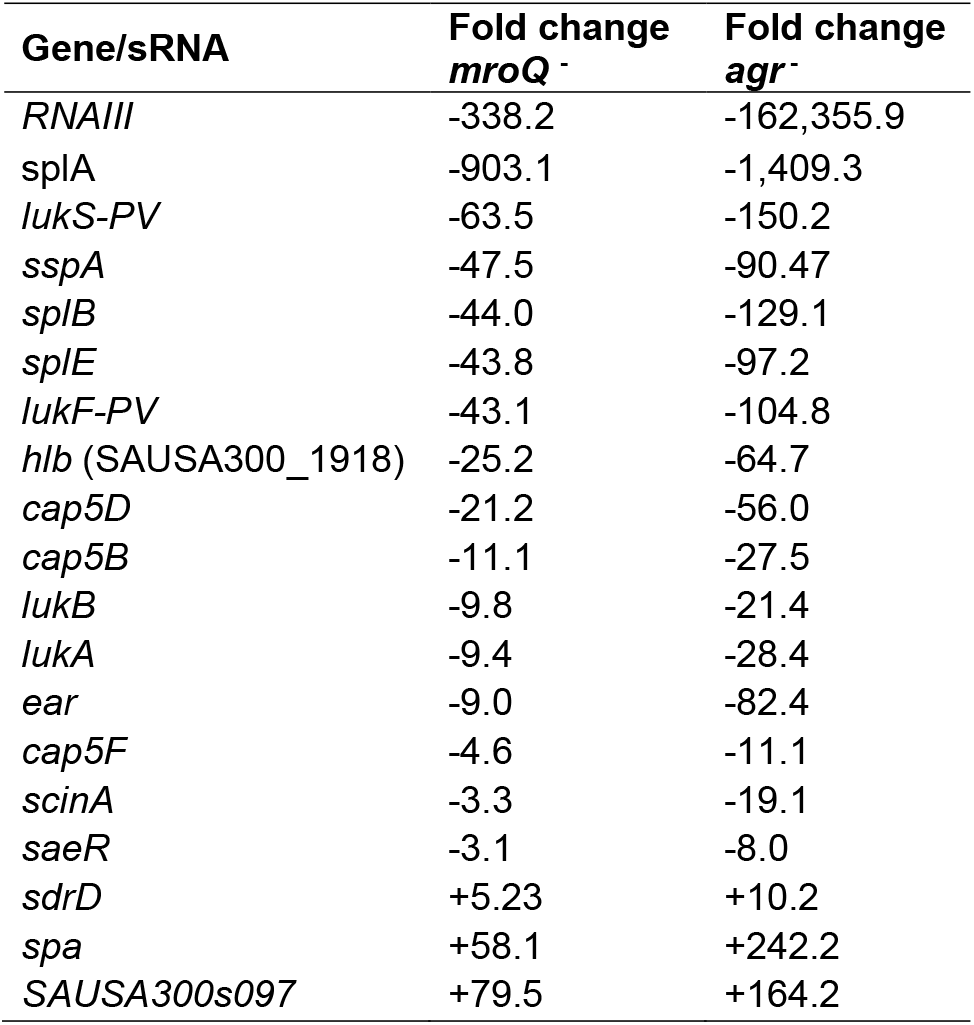
Comparison of genes/sRNA demonstrating changes in expression in both *mroQ*^−^ and *agr*^−^ mutant RNA-seq datasets.

### Complementation of the *mroQ* mutation

A *mroQ* complementing PCR fragment was created using OL4139 and OL4140 (all primers used are listed in Table 2), with the former being located immediately at the *mroQ* start codon and the latter 88 bp downstream of the 3’ end of the coding region. We also performed site-directed mutagenesis to create a catalytically inert complementing fragment, using splicing by overhang extension (SOE) PCR techniques as previously described by us (60). This was generated using primers OL4139/OL3613 and OL3612/OL4140, where OL3612 and OL3613 contained mutated nucleotide sequences that change the native ‘GAAGAA’ sequence encoding for two glutamic acid residues to a ‘GCAGCA’ sequence encoding for two alanine residues. To facilitate Western blot analysis a hexahistidine (6x-His) tag was added to primer OL4140, creating OL3777, and used along with OL4139 to create a tagged fragment for localization studies. Each of these three *mroQ* fragments were separately cloned into pJB67, a multi-copy plasmid with a CdCl2 inducible promoter and ribosome binding site (61). An additional *mroQ* complementing strain was created using OL4332 and OL4333, which are identical to OL4139 and OL4140 with the exception of differing restriction sites that were compatible with pEPSA5, a xylose-inducible shuttle vector.

All constructs were transformed into chemically competent *E. coli* (DH5α), with clones confirmed using gene specific primers, followed by Sanger sequencing (Eurofins MWG Operon). Plasmids bearing the correct sequence were transformed into *S. aureus* RN4220 by electroporation and confirmed by PCR. Following this, a phage lysate was created using Φ11, which was subsequently used to transduce the *S. aureus* USA300 LAC *mroQ* mutant. Strains were confirmed via Sanger sequencing using plasmid specific primers. Additionally, the pJB67 empty vector was transformed into *S. aureus* RN4220 by electroporation and confirmed by PCR. A phage lysate was created using Φ11, which was then used to transduce the *S. aureus* USA300 LAC wild type, USA300 LAC *mroQ* mutant, USA300 LAC *agr* mutant, and USA300 LAC *sigB* mutant, to serve as negative controls. Strains were subsequently confirmed via PCR with plasmid specific primers OL4093 and OL4094. These control strains were also created with the pEPSA5 empty vector, as described above, and confirmed via PCR with plasmid specific primers OL4358 and OL4359.

### Construction of *lacZ*-reporter fusion strains

*hla-lacZ* and *aur-lacZ* fusion strains were previously constructed using plasmid pAZ106 (62, 63). Phage lysates were created from SH1000 pAZ106::*aur-lacZ* and SH1000 pAZ106::*hla-lacZ* strains using Φ11, before transducing both separately into the USA300 LAC wild type, alongside its *mroQ* and *agr* mutant strains. Following this, strains containing fusions were confirmed via PCR using a reverse primer located downstream of the MCS and within the *lacZ* gene, OL1232, and the fusion’s respective forward primer.

### β-galactosidase assays

Levels of β-galactosidase activity were quantified for *aur-lacZ* and *hla-lacZ* fusions as described previously (62). Assays were performed in biological triplicate and technical duplicate. Strains were grown as described above and standardized to an OD_600_ of 0.05. Samples were taken every hour for 6 h and processed using 4-Methylumbelliferyl beta-D-Galactopyranoside (4-MUG) as a substrate. A Student’s t-test with Welch’s correction was used to determine statistical significance.

### Zymogram and western blot analysis

Zymograms were performed as previously described (62), using culture supernatants from strains grown for 15 h. The OD_600_ of these cultures were standardized to the lowest value amongst the collection, before being concentrated using Amicon Ultra-2 centrifugal 3 kDa filter units (Millipore Sigma). For Western blot analysis, overnight bacterial cultures were grown in biological triplicate as described above, standardized to an OD_600_ of 0.05 and grown to appropriate time points. Secreted (5 h) and insoluble (3 h) protein fractions were harvested as previously described (64) before being run on 12% SDS-PAGE gels. Samples were subsequently blotted to PVDF membranes as previously described (33). Immunoblotting was performed with anti-Protein A mouse monoclonal primary antibody (SPA-27, Sigma Aldrich) or anti-LukS-PV mouse monoclonal primary antibody (IBT), overnight at 4°C. The secondary antibody used was HRP-conjugated goat anti-mouse IgG (Cell Signaling Technologies). For localization studies, the *mroQ*-6xHis complement strain was grown overnight in biological triplicate as described above. Next, 1 ml of bacterial culture was centrifuged, and supernatants were collected and concentrated overnight using 10% trichloroacetic acid (TCA); with the pellet processed as previously described (64) to harvest the various protein fractions (cytoplasmic and membrane). Samples were prepared for immunoblotting as described above, and immunoblotting was performed using anti-6xHis oligoclonal rabbit primary antibody (ThermoFisher Scientific) overnight at 4°C. The secondary antibody used was HRP-conjugated goat anti-rabbit IgG (Cell Signaling Technologies). HRP activity was assessed using the SuperSignal^™^ West Pico Chemiluminescent substrate (ThermoFisher Scientific) and visualized on X-ray film.

### Hemolysis assay

Overnight bacterial cultures were grown in biological triplicate as described above and standardized to an OD_600_ of 0.05. Cultures were incubated for 6h before being centrifuged, and the supernatant removed and mixed (1:1) with hemolysin buffer (0.145 M NaCl; 20 mM CaCl_2_), followed by the addition of 25 μl whole human blood (BioIVT). Samples were incubated for 40 min at 37°C on a rotator, before being centrifuged at 5.5 x g for 1 min. Supernatants were collected and hemolysis was determined by measuring the OD_543_ using a BioTek Synergy II plate reader and 96-well plates. A Student’s t-test with Welch’s correction was used to determine statistical significance.

### qPCR analysis

Quantitative real-time PCR (qPCR) was performed on samples isolated at hour 5 in biological triplicate as previously described (65). Various targets were examined and normalized to 16s rRNA (primers are listed in Table 2). A Student’s t-test with Welch’s correction was used to determine statistical significance of expression units.

### Transcriptomic analysis via RNA-sequencing

RNA-seq was performed as previously described (66). Briefly, USA300 LAC wild type, alongside its *mroQ* and *agr* mutant strains, were standardized and grown in biological triplicate for 5 h using the growth method described above. Next, 5 ml of each culture was immediately combined with 5 ml of ice-cold PBS, pelleted by centrifugation at 4°C, and the supernatant removed. Total RNA was isolated using an RNeasy kit (Qiagen), with DNA removed using a TURBO DNA-free kit (Ambion). DNA removal was confirmed by PCR using OL398 and OL399, and RNA quality was assessed using an Agilent 2100 Bioanalyzer and an RNA 6000 Nano kit (Agilent) to confirm integrity; samples with a RIN greater than 9.7 were used. Biological replicates were pooled at equal RNA concentrations, before rRNA depletion using a Ribo-Zero Kit for Gram-Positive Bacteria (Illumina), and MICROBExpress Bacterial mRNA enrichment kit (Ambion). Removal efficiency was confirmed, again using an Agilent 2100 Bioanalyzer and RNA 6000 Nano kit. Strand-specific library preparation was performed using a total RNA-seq Kit v2 and whole-transcriptome protocol (Ion Torrent). Sample libraries were assessed for quality, concentration, and average fragment size using the Agilent 2100 Bioanalyzer and High Sensitivity DNA kit (Agilent). Prepared fragment libraries were ligated to Ion Sphere Particles, amplified, and enriched using an Ion Torrent Personal Genome Machine OT2 200 Kit and Ion OneTouch 2 System. Particles were then sequenced using an Ion 318 v2 chip (Ion Torrent).

### Bioinformatics

Raw data files (.fasta) were exported from the Ion Torrent PGM Server using the File Exporter plugin and uploaded to Qiagen Bioinformatics Workbench for analysis. Reads corresponding to ribosomal RNA were filtered, removed by aligning to known rRNA sequences, and discarded. Unmapped (filtered) read sequences were aligned to the USA300 FPR3757 reference genome (NC_007793.1) and expression values were calculated using the RNA-Seq Analysis function. Experimental comparisons were carried out following quantile normalization through the RNA-seq experimental fold change feature. Expression values calculated for each gene are shown as normalized RPKM values. Experimental data from this study was deposited into the NCBI Gene Expression Omnibus (GEO accession: GSE121625). Reviewer Link: https://www.ncbi.nlm.nih.gov/geo/query/acc.cgi?acc=GSE121625 (Token: spqxgoskftcvvwd).

### Biofilm formation assay

The ability to form biofilms was examined for our strains using methods previously described (33). Overnight bacterial cultures were grown in biological triplicate as described above and standardized to an OD_600_ of 0.5. Cultures were then seeded in a 24-well plate containing TSB, 5% human plasma, and erythromycin for plasmid retention, in technical duplicate, and grown for 24 h. Following incubation, biofilms were washed with PBS, and briefly fixed with 100% ethanol. Biofilms were then allowed to dry overnight prior to staining with crystal violet for 15 min. Following this, wells were washed to remove excess crystal violet, and allowed to dry overnight. Biofilm formation was quantified by extracting crystal violet stain with 100% ethanol, and measuring OD_550_ using a BioTek Synergy II plate reader. A Student’s t-test with Welch’s correction was used to determine statistical significance.

### Oxidative stress assay

Overnight bacterial cultures were grown in biological triplicate as described above and standardized to an OD_600_ of 0.1. Samples were treated with 20 mM hydrogen peroxide for 30 min with shaking at 37°C. After this time, the reaction was stopped by the addition of 10 μg/ml catalase. Percent recovery was determined by comparing pre-exposure CFU/ml to post-exposure CFU/ml. A Student’s t-test with Welch’s correction was used to determine statistical significance.

### Murine model of sepsis and dissemination

These experiments were performed as previously described (67). Briefly, six-week old, female CD-1 mice were purchased from Charles River Laboratories and allowed to acclimate for one week prior to the start of experimentation. Wild type and mutant strains were grown as described above and standardized to an OD_600_ of 0.05. Cultures were grown overnight for 15 h on three separate days, and the average CFU/ml across replicates was calculated for each strain. This average CFU/ml was used to determine the volume of bacteria needed to obtain a 10 ml inocula of 5 x 10^8^ ((5 x 10^8^ / average CFU/ml) x 10 ml). On the day of infection, appropriate aliquots from fresh overnight cultures prepared in the same way were centrifuged, resuspended in 1 ml sterile PBS to wash, centrifuged once more, and finally resuspended in a total volume of 10 ml sterile PBS. Next, 100 μl of bacterial suspension was administered via tail vein injection, providing a final inocula of 5 x 10^7^ CFU/ml. Infections were monitored over a 7 day period, or until mice reached a premoribund state, at which point they were euthanized (67). At this time, kidneys were harvested and stored at −80°C. These were subsequently photographed before being homogenized in 1 ml PBS, and serial diluted onto TSA to determine bacterial burden (CFU/ml). A Mann-Whitney test was performed to determine statistical significance for bacterial burden in kidneys.

### Murine model of skin abscess formation

These experiments were performed as described previously (68). Briefly, wild type and mutant strains were grown for 2.5 h until reaching an OD_600_ of approximately 0.75. Bacterial cells were centrifuged, and the pellet was resuspended in sterile PBS for inoculation. Inocula were prepared at 1 x 10^7^ CFU / 50 μl for injections, and subsequently confirmed via serial dilution and plating. Six-week old BALB/c female mice were shaved on the right flank prior to treatment with Nair for full hair removal. Subsequently, mice were injected in the right flank with 50 μl and infections were allowed to persist for 7 days. Throughout the infection period, mice were monitored for irregular activity or distress. Following the 7 day infection period, mice were euthanized, and abscesses were excised, weighed, and measured (length and width) to determine abscess area (mm^2^). Abscesses were then homogenized, and homogenates were diluted and plated onto TSA to determine bacteria burden (CFU/g). A Student’s t-test with Welch’s correction was used to determine statistical significance.

### Mass spectrometry of culture supernatants for AIP-I detection

The wild type, *mroQ* mutant, *agr* mutant, and *mroQ*-pEPSA5 complement were grown for 15 hours in TSB supplemented with 1% xylose as described above. Cultures were pelleted, and supernatants were filter sterilized (0.2 μm) to ensure they were cell free. Samples were then processed by the Triad Mass Spectrometry Facility at The University of North Carolina at Greensboro, alongside controls of TSB and purified AIP-I (AnaSpec Inc.; Fremont CA). Samples were analyzed using an Acquity UPLC (Waters, Milford, MA) coupled to a TSQ Quantum Access Triple Quadrupole mass spectrometer (Thermo, San Jose, CA) with heated electrospray ionization (HESI) source. A 5 μL injection of each sample was eluted from an Acquity BEH C18 column (2.1 mm × 50 mm, 1.8 μm packing) using a binary solvent system consisting of optima grade water and optima grade acetonitrile (both containing 0.1% formic acid) at a flow rate of 300 μL/min. The gradient initiated at 75:25 water:acetonitrile and increased linearly to 50:50 water:acetonitrile over 3.5 minutes. The column was then washed with 100% acetonitrile and reequilibrated to starting conditions. The mass spectrometer was operated in MRM mode using positive ion electrospray ionization. The following transitions were monitored: *m/z* 961.3 ([M+H]^+^) to *m/z* 711.2 and *m/z* 961.3 to *m/z* 365.9 at a gas pressure of 1.5 and collision energy of 29 and 43, respectively. Quantitative analysis was performed using the *m/z* 961.3 to 711.2 transition. The HESI source was operated using the following parameters: tube lens offset, 198 V, spray voltage, 5.00 kV, sheath gas pressure, 60, auxiliary gas pressure, 35, capillary temperature, 200°C, and capillary offset, 35 V.

## Supplemental Material

Supplemental material for this article may be found at (link)

Figure S1-S5, Table S1-S5, PDF file

## Acknowledgements

This work was supported in part by AI080626 and AI124458 (both LNS) from the National Institute of Allergy and Infectious Diseases, and by AT006860 from the National Center for Complementary and Integrative Health (to NBC and DAT).

